# Characteristics and dynamical signatures of recurrent cortical circuits during context-dependent processing

**DOI:** 10.64898/2026.02.06.704473

**Authors:** Yue Kris Wu, Ho Yin Chau, Serena Di Santo, Kenneth D. Miller

## Abstract

Context profoundly shapes neural responses and behavior. During context-dependent sensory processing, recurrent connections shape the integration of feedforward sensory input and feed-back input from downstream brain regions. How do different cell types, interacting through spatially structured recurrent lateral connections, give rise to context-dependent processing and circuit stability, and what dynamical signatures reveal their individual roles? To answer these questions, we employ data-driven approaches to construct spatially extended stabilized supralinear network models that capture the responses of diverse cell types in the mouse primary visual cortex during context-dependent processing. Analysis of well-fitting models reveals that the dominant inhibitory cell type affecting excitatory neurons is not fixed but dynamically varies with stimulus and space. While PV-mediated stabilization is indispensable across all models and stimulus conditions, SST-mediated stabilization is also required, and likely in a stimulus-dependent manner. Interestingly, even when a specific inhibitory cell type is required for circuit stabilization, a uniform perturbation of it does not necessarily produce a paradoxical change in its mean activity. Instead, assessing cell-type-specific circuit stabilization requires patterned perturbations, where paradoxical effects manifest along specific activity modes. Finally, we show that recurrent connections and input-output nonlinearities are essential for integrating feedforward and feedback inputs to reproduce the observed spatial response profiles. Recurrent excitatory connections, in particular, are required to account for responses to small stimuli, where external inputs are relatively weak. Taken together, our work reveals the crucial role of ubiquitous biological components in context-dependent processing and delineates the characteristics and dynamical signatures of these circuits.

## Introduction

Our perception of a sensory stimulus is profoundly influenced by the context in which the stimulus is presented (Bar, 2004; Kuchibhotla et al., 2017). For example, optical illusions can arise when the color, light, and patterns in the visual scene are arranged in specific ways (Eagleman, 2001). Similarly, sound at a given volume may be perceived as louder in a quieter environment (Moore, 1977). Understanding context-dependent sensory processing, a fundamental computation to extract meaning from the environment, is essential for comprehending sensory perception.

In the visual system, contextual information can be derived from the visual scene surrounding the stimulus. Conventionally, excitatory neurons in the primary visual cortex (V1) are thought to be driven primarily by stimuli confined to a limited region of visual space, known as the classical receptive field (Hubel and Wiesel, 1959). However, neural responses are strongly modulated by context (Carandini and Heeger, 2012). For example, increasing stimulus size beyond the classical receptive field typically reduces the response of neurons whose receptive fields are centered on the stimulus, a phenomenon referred to as *surround suppression* (DeAngelis et al., 1994; Li and Li, 1994; Cavanaugh et al., 2002; Ozeki et al., 2009). This strong contextual modulation of neural activity is not restricted to excitatory neurons but is also observed across multiple inhibitory cell types, including parvalbumin-expressing (PV), somatostatin-expressing (SST), and vasoactive intestinal peptide-expressing (VIP) interneurons (Adesnik et al., 2012; Keller et al., 2020a).

These context-dependent effects on neural activity arise from the combined contributions of feedforward and feedback inputs, as well as recurrent interactions. Recent work has examined how distinct input sources drive neural activity (Di Santo et al., 2025), however, little is known about how recurrent connections shape the integration of feedforward and feedback signals during contextdependent processing, and about the characteristics and dynamical signatures that underlie these recurrent circuits. A major challenge is disentangling the complex interplay among diverse cell types and the recurrent lateral interactions that unfold across space. To address this challenge, we employ data-driven modelling approaches grounded in biologically realistic circuit models and tightly constrained by experimental measurements. Specifically, we infer spatially extended stabilized supralinear network models that are fitted to the responses of multiple cell types in mouse V1 during context-dependent processing.

Analysis of well-fitting models reveals several key features of the underlying recurrent circuits. First, the inhibitory cell type exerting the strongest influence on excitatory neurons is not fixed, but instead varies dynamically with stimulus conditions and spatial location. Second, the well-fitted models operate in an inhibition-stabilized regime. While PV-mediated stabilization is indispensable across all stimulus conditions, SST interneurons also play a critical and likely stimulus-dependent role in stabilization. Importantly, we find that uniform perturbations of an inhibitory cell type required for stabilization do not necessarily induce a paradoxical change in its mean activity.

Instead, uncovering cell-type-specific stabilization requires patterned perturbations, which reveal paradoxical effects along specific activity modes rather than in global averages. Finally, we show that recurrent connectivity and input-output nonlinearities are essential for integrating feedforward and feedback inputs to reproduce observed spatial response profiles. Recurrent excitatory connections, in particular, are crucial for explaining responses to small stimuli, where external inputs are relatively weak and recurrent amplification becomes necessary.

Taken together, our results highlight how ubiquitous biological components, such as recurrent connectivity and nonlinear input-output transformations, jointly enable context-dependent processing. By delineating the characteristic dynamical signatures of these circuits, our work provides insight into the interpretation of perturbation experiments and the properties of the recurrent circuits that implement context-dependent processing.

## Results

To elucidate how spatially distributed recurrent interactions shape context-dependent processing, we apply data-driven approaches to construct spatially extended stabilized supralinear network models constrained by experimental measurements. Specifically, we consider responses of V1 layer 2/3 (L2/3) neurons in awake, head-fixed mice to visual stimuli of varying sizes (Fig. 1). In our modeling setup, the L2/3 recurrent cell types considered include E, PV, SST, and VIP. These populations receive feedforward excitatory input from layer 4 (L4), feedback excitatory input from the lateromedial visual area (LM), a higher visual cortical area that provides prominent feedback projections to V1 (Wang and Burkhalter, 2007; Marques et al., 2018), as well as a residual input (R) representing contributions from other brain regions to V1 L2/3 (Fig. 1). Neurons with different receptive fields are modeled as units positioned at different locations in a two-dimensional retinotopic coordinate space. In a continuous formulation, the rate dynamics of neurons of L2/3 recurrent cell type *A* located at position **x** in response to stimulus size *s* is given by:

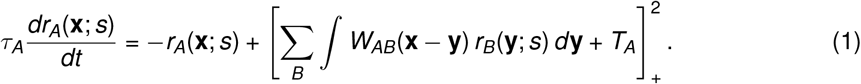

**Fig. 1.**
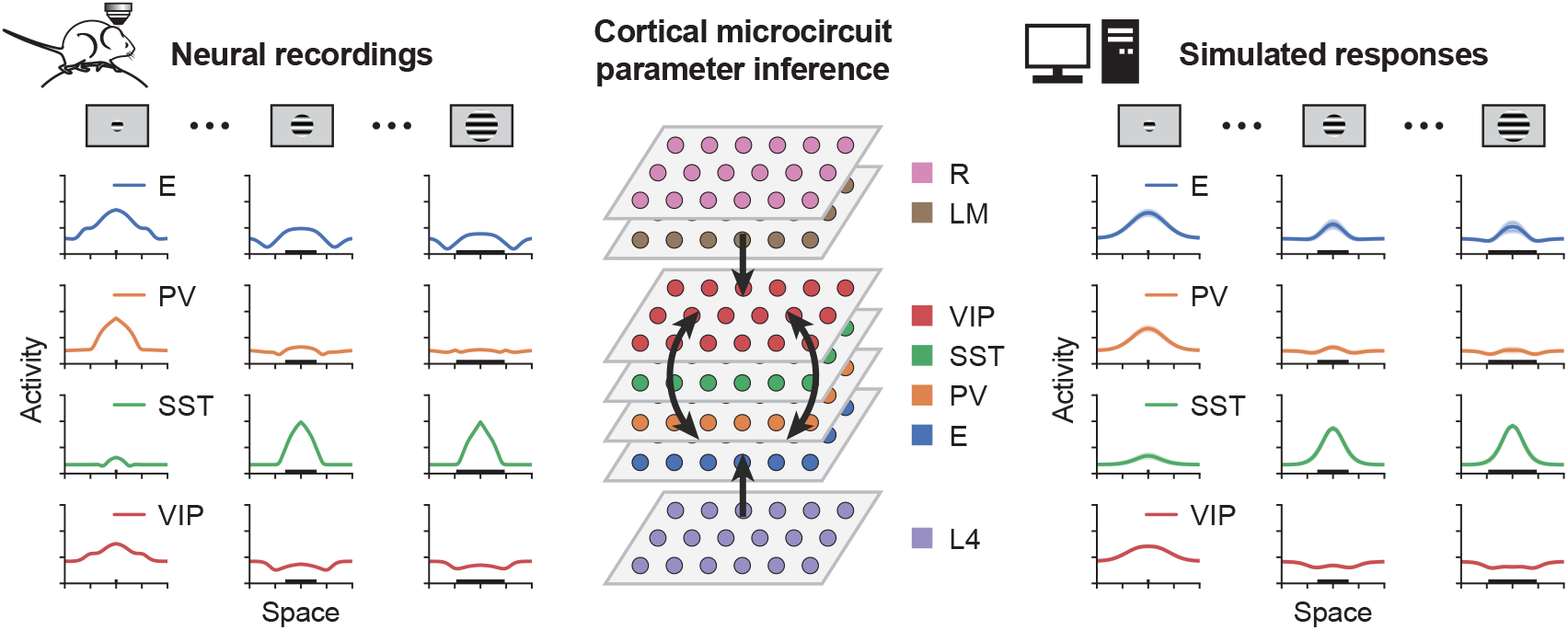
Spatially extended cortical microcircuit parameter inference pipeline. Awake, head-fixed mice were presented with drifting grating stimuli of varying sizes, and neural activity was recorded using two-photon calcium imaging from E, PV, SST, and VIP neurons in V1 L2/3, as well as from excitatory neurons in V1 L4 and excitatory boutons originating from lateral medial cortex (LM) within V1 (Keller et al., 2020a,b). Responses from neurons with different receptive fields were quantified in the retinotopic coordinate space. These experimentally measured response profiles were then used to infer the parameters of spatially extended cortical microcircuit models. Model parameter optimization was performed using backpropagation through time to minimize error between simulated and experimentally measured responses of the recurrent cell types. The best-fitting models were subsequently analyzed in detail to elucidate the characteristics and dynamical signatures of recurrent cortical circuits during context-dependent processing.

Here, *τ*_*A*_ is the time constant of the rate dynamics. *W*_*AB*_(**x**−**y**) denotes the connection strength from neurons of cell type *B* at location **y** to neurons of cell type *A* at location **x**. *T*_*A*_ is the bias, and 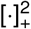 denotes a rectified quadratic function that captures the experimentally measured input-output relation (Hansel and van Vreeswijk, 2002; Miller and Troyer, 2002; Priebe et al., 2004; Ahmadian and Miller, 2021). Cell types are defined as *A* ∈ {*E, PV, SST, VIP*} and *B* ∈ {*E, PV, SST, VIP, L*4, *LM, R*}. The feedforward input from L4 and the feedback input from LM are determined by experimental data (Fig. S1, Keller et al., 2020a,b; Di Santo et al., 2025), while the residual input is modeled as a simple two-dimensional Gaussian with fixed spatial scale and a peak amplitude that increases with stimulus size (Fig. S2, see Methods). The spatial profiles of the connectivity kernels *W* are modeled as two-dimensional Gaussians, with spatial scales extracted from previous literature (Di Santo et al., 2025). The connection strengths (i.e., amplitudes of *W*) and bias terms *T* are optimized using backpropagation through time to minimize error between simulated and experimentally measured responses of the recurrent cell types. (Fig. 1, see Methods). Following optimization, we analyze the top ten models with the lowest errors, which recapitulate the data and consistently show that excitatory neurons with receptive fields centered on the stimulus elicit the largest responses across space for all stimulus sizes (Fig. S1).

### Context shifts the dominance of cell-type-specific inhibition to the excitatory population

The resulting best-fit models well reproduce the experimentally measured rate fields, which quantify how neurons with receptive fields at different spatial locations respond to a given stimulus (Fig. 2A, Fig. S3). In addition, the models closely capture the experimentally measured size-tuning curves, which describe how neurons with receptive fields centered at the stimulus center (i.e., at 0° in retinotopic space) change their responses with stimulus size (Fig. 2B). Interestingly, although connection strengths are unconstrained during optimization, the resulting models naturally recapitulate experimentally observed connectivity features, such as negligible connection strengths from SST to SST and from VIP to VIP, and weak connection strengths from VIP to E and PV (Pfeffer et al., 2013), as shown in Fig. S4. To identify the characteristics of these well-fitted models, we first analyze the spatial input currents from different cell types to excitatory neurons across stimulus conditions. We find that in all the models, across stimuli and space, recurrent excitation provides the dominant source of excitation to excitatory neurons (Fig. 3A) and the other L2/3 cell types (Fig. S5). Strikingly, for small stimuli, across space, PV provides the dominant inhibition to excitatory neurons (Fig. 3A left, 3B). For larger stimuli, however, SST dominates inhibition onto excitatory neurons at the center, while PV exerts stronger inhibition in the surround (Fig. 3A right, 3B). These results indicate that the dominance of cell-type-specific inhibition onto excitatory neurons is not fixed but flexibly reconfigures with context. Furthermore, the overall current provided by the residual input is relatively weak compared to L4 and LM (Fig. 3A, Fig. S5, Fig. S6). Note that despite being relatively weak, the residual input is important for generating the appropriate excitatory activity and maintaining high SST activity at large stimulus sizes (Fig. S7, Fig. S8).

**Fig. 2.**
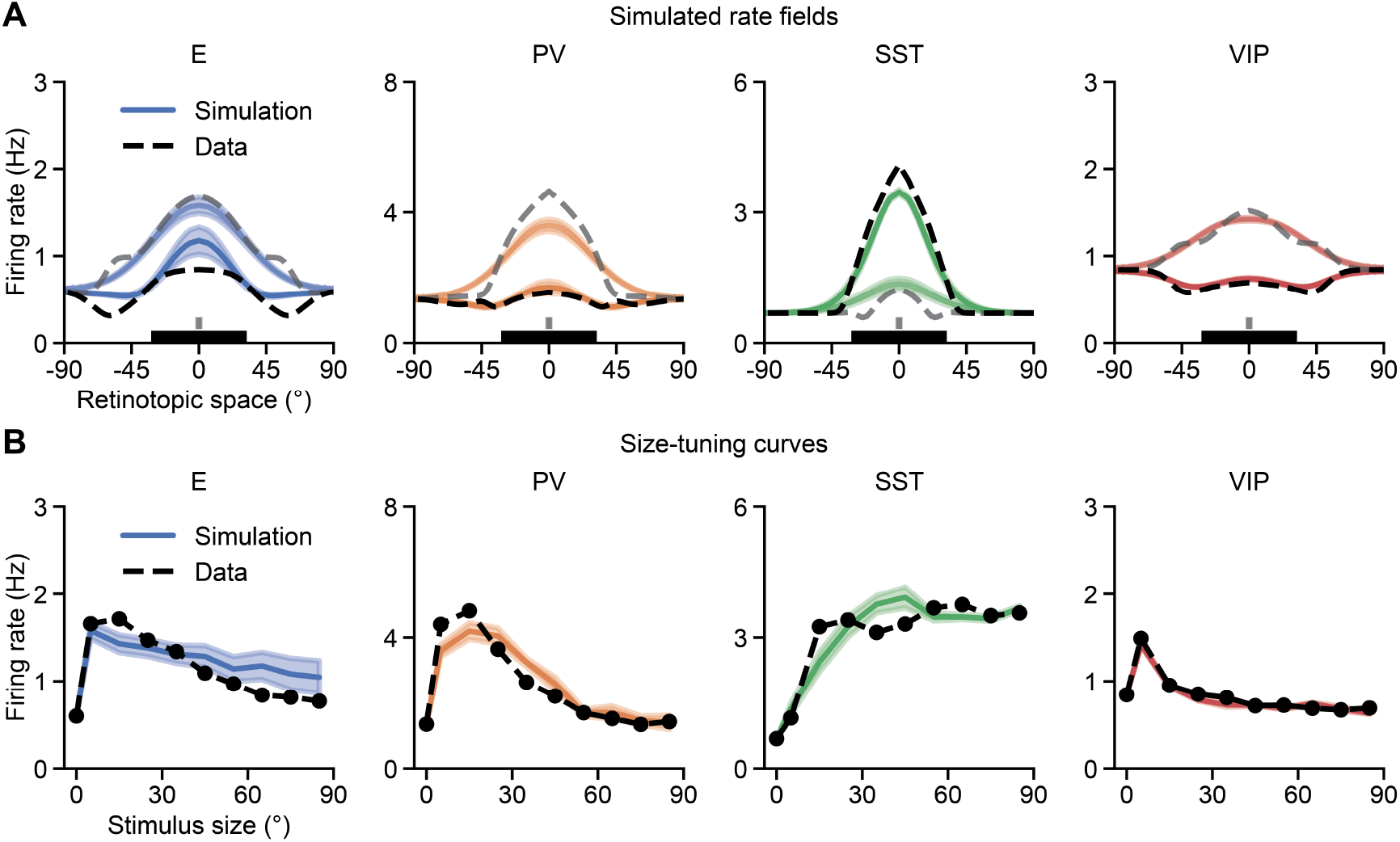
**A**. Comparison of simulated rate fields from optimized models (colored) and experimental data (black) for E, PV, SST, and VIP populations across space for two example stimulus sizes (5^*°*^and 65^*°*^), shown as lines of varying brightness. The lighter lines correspond to 5^*°*^, and the darker lines correspond to 65^*°*^. Shaded regions indicate the standard deviation of the corresponding rate fields from the top ten optimized models. Bars of different lengths at the bottom indicate the two example stimulus sizes. **B**. Comparison of model and data size-tuning curves for E, PV, SST, and VIP populations. Size-tuning curves are quantified using the responses of neurons from each cell type located at 0^*°*^in retinotopic space. Shaded regions indicate the standard deviation of the corresponding size-tuning curves.

**Fig. 3.**
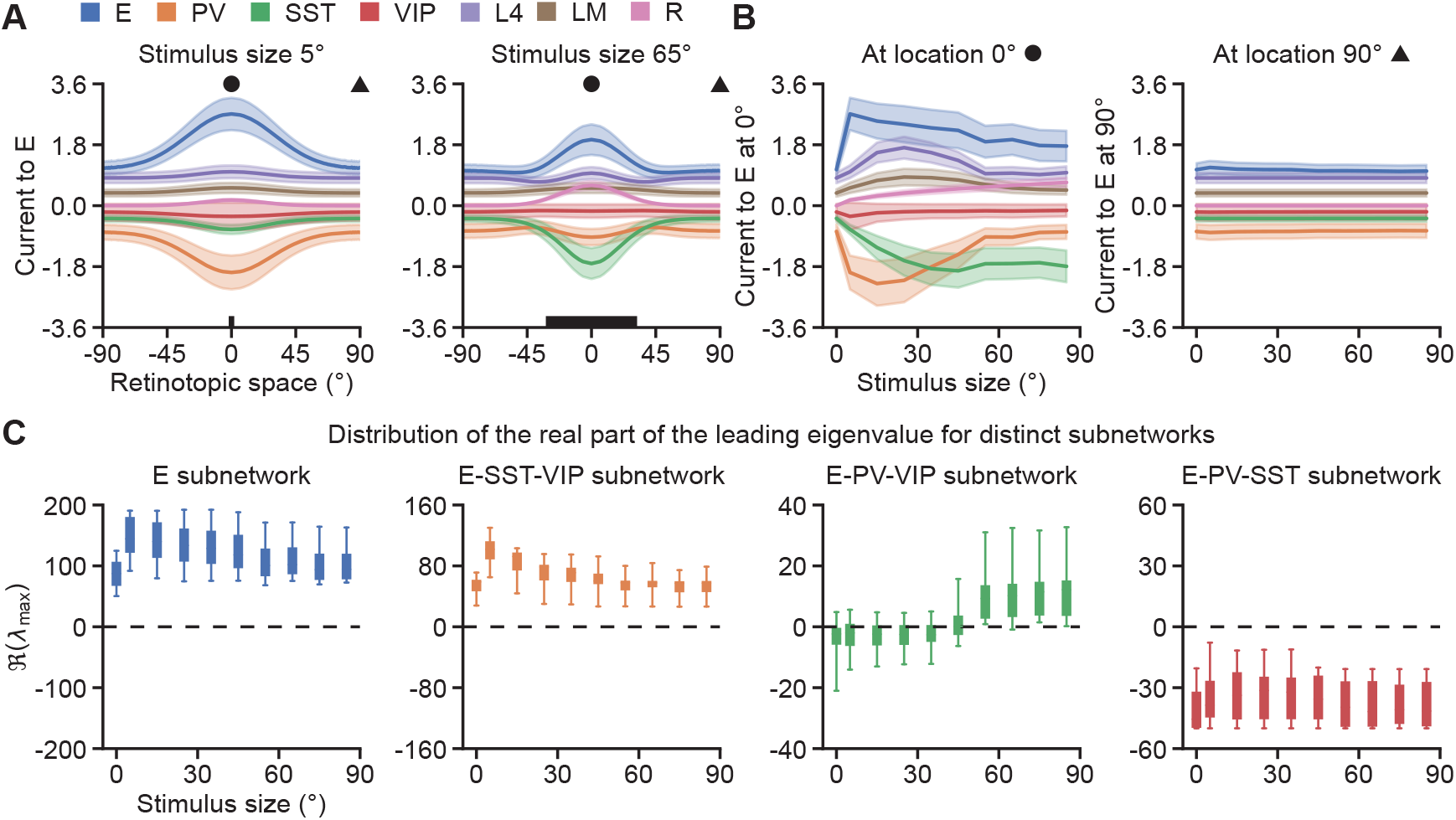
**A**. Left: Current from different sources to the E population at stimulus size 5°. Solid lines and shaded regions indicate the mean and the standard deviation of the corresponding currents, respectively. Right: Same as left but for stimulus size 65°. **B**. Left: Current from different sources to the E neurons located at 0° in the retinotopic space (marked by a circle in A) as a function of stimulus size. Right: Same as left but for the E neurons located at 90° in the retinotopic space (marked by a triangle in A). **C**. Distribution of the real part of the leading eigenvalue of the Jacobian of different subnetworks across stimulus sizes. From left to right: E subnetwork, E-SST-VIP subnetwork, E-PV-VIP subnetwork, and E-PV-SST network.

### Context affects cell-type-specific inhibition stabilization properties

Given the context-dependent shift in the dominance of cell-type-specific inhibition, we wonder whether this shift could imply a corresponding change in cell-type-specific inhibition stabilization, that is, a change in the requirement of a particular inhibitory cell type for stabilization. To this end, we analyze the eigenspectrum of the Jacobian of the subnetwork in which a given inhibitory cell type is excluded across stimulus sizes. A positive (negative) real part of the leading eigenvalue of that Jacobian indicates that the excluded cell type or types are required (not required) for stabilization (see Methods). We find that, across all models and stimulus sizes, the real part of the leading eigenvalue is positive for both the E subnetwork and the E-SST-VIP subnetwork, but negative for the E-PV-SST subnetwork (Fig. 3C). These results indicate that all model networks are inhibition-stabilized, with PV-mediated stabilization being indispensable, whereas VIP neurons do not contribute to stabilization. In contrast to those subnetworks, the E-PV-VIP subnetwork exhibits a moderate degree of heterogeneity, highlighting both circuit degeneracy and inter-individual variability. Specifically, although a small subset of models show a positive real part of the leading eigenvalue across all stimulus sizes, most models exhibit this only for large stimuli (Fig. S9). And the mean of the leading eigenvalue distribution gradually shifts from negative to positive as stimulus size increases (Fig. 3C), suggesting that SST-mediated stabilization is also required and likely in a stimulus-dependent manner.

### Patterned, but not uniform, perturbations reveal cell-type-specific inhibitory stabilization

Given the challenges of directly estimating eigenspectra from experimental recordings, we aim to identify dynamical signatures that can reflect the characteristic features of the eigenspectra. Inhibition stabilization is commonly associated with paradoxical effects, whereby inhibitory activity paradoxically decreases in response to excitatory current injection into the inhibitory population (Tsodyks et al., 1997; Ozeki et al., 2009; Sanzeni et al., 2020; Miller and Palmigiano, 2020; Sadeh and Clopath, 2021). Most previous computational studies have focused on either population models or networks lacking explicit spatial structure (Litwin-Kumar et al., 2016; Molino et al., 2017; Mahrach et al., 2020; Palmigiano et al., 2020; Waitzmann et al., 2024; Bos et al., 2025). In these studies, perturbations are typically applied uniformly to the inhibitory population, that is, every inhibitory neuron receives the same perturbative current. To examine the consequences of such perturbations in spatially extended networks, we then apply uniform excitatory perturbations to all inhibitory populations, or selectively to PV, SST, or VIP populations across space, to a network responding to a stimulus of size 55°. Note that, in all models, SST is required for network stabilization given a 55°stimulus (Fig. 3C, Fig. S9). Across all models, uniformly perturbing all inhibitory populations paradoxically reduces the mean inhibitory activity (Fig. 4A). Similarly, uniform perturbations to PV populations paradoxically decrease the average PV activity, whereas uniform perturbations to VIP populations increase the average VIP activity (Fig. 4A). In contrast, the same perturbations of SST populations yield distinct response patterns in different models (Fig. 4A). That is, even though SST is required for circuit stabilization, a uniform perturbation of SST does not necessarily induce a paradoxical change in its mean activity (Fig. 4A).

**Fig. 4.**
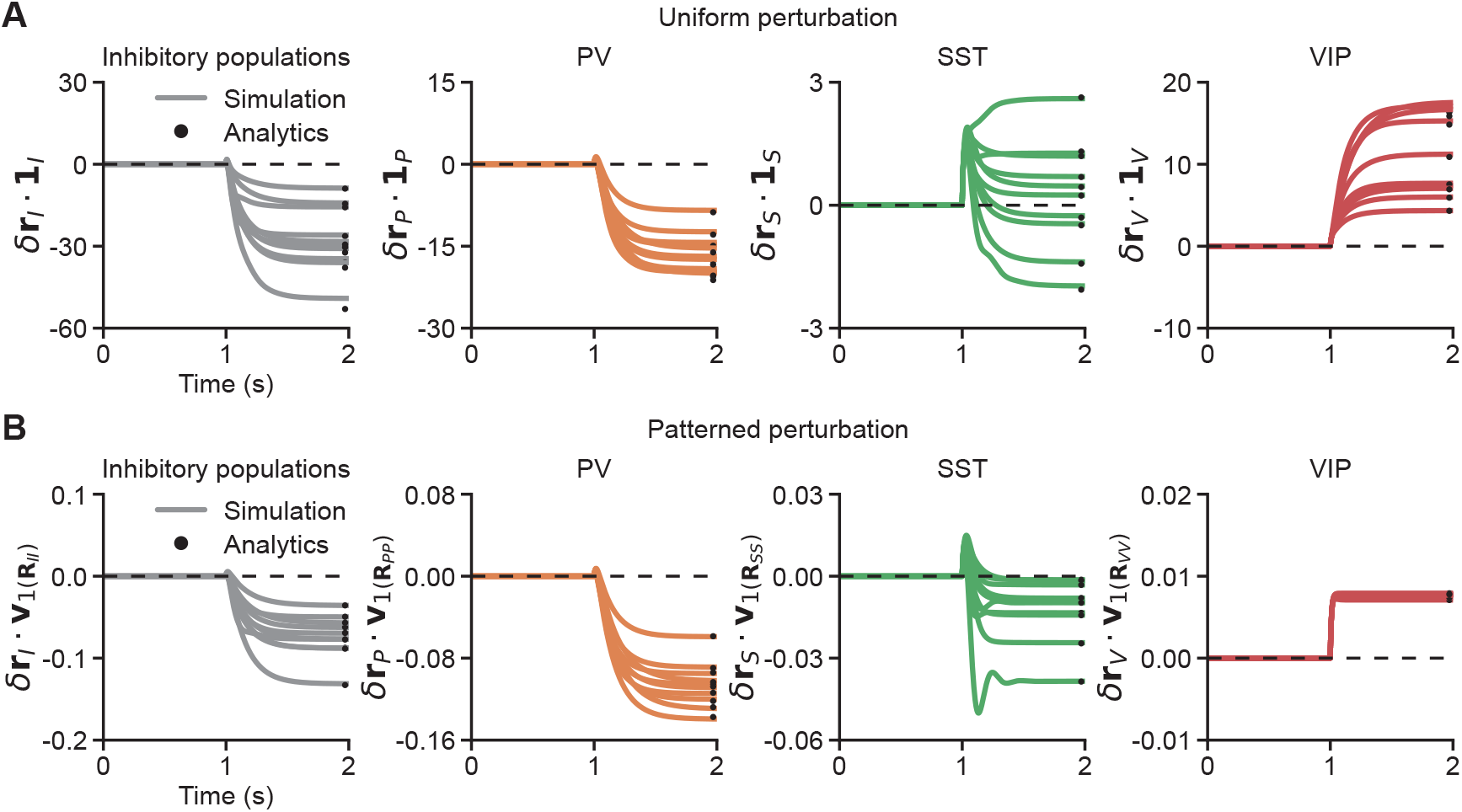
**A**. From left to right: starting from the steady-state response to a sustained 55°stimulus, measurements of the changes in the mean activity of inhibitory, PV, SST, and VIP populations induced by uniform excitatory perturbations, quantified as the dot product between cell-type-specific activity changes and an all-ones vector. Uniform excitatory perturbations (*γ***1**_*X*_ with *γ* = 0.001, where **1**_*X*_ is a vector of all 1’s for population *X*) are applied from 1 s onward to all inhibitory, PV, SST, or VIP neurons, respectively. For a 55°stimulus, SST is required for stabilization across all models (Fig. 3B, Fig. S9). Each line corresponds to the simulation result from a single model. Black dots indicate analytical results calculated using Eq. 5. The top ten models are shown. **B**. Same as A, but for patterned perturbations, quantified as the dot product between cell-type-specific activity changes and the patterned perturbation vector. Patterned perturbations (*γ***v**_1_ with *γ* = 0.005, where the eigenvector **v**_1_ is associated with the smallest eigenvalue of the corresponding response matrix **R**_*XX*_, Eq. 4 and Eq. 6) are applied from 1 s onward to all inhibitory, PV, SST, or VIP neurons, respectively, at stimulus size 55°, where SST is required for stabilization across all models. Black dots indicate analytical results calculated using Eq. 6. The top ten models are shown.

To understand why uniformly perturbing a cell type required for stabilization does not necessarily induce a paradoxical change in its mean activity, we apply linear response theory to examine how changes in the activity of cell type *X, δ***r**_*X*_, depend on changes in their input, *δ***h**_*X*_, and express this relationship as

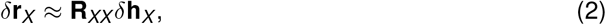

where **R**_*XX*_ is the block diagonal matrix of the response matrix **R** (see Methods) that specifies how a perturbative input to cell type *X* is transformed into changes in the activity of cell type *X*. A uniform excitatory perturbation to cell type *X* can be written as *γ***1**_*X*_ where *γ >* 0 controls the perturbation amplitude and **1**_*X*_ denotes the all-ones vector. The vector **1**_*X*_ can be expressed as a linear combination of the eigenvectors of **R**_*XX*_ :

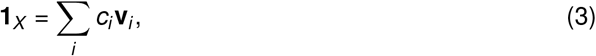

with

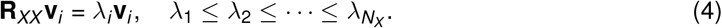

Here, *c*_*i*_ are the coefficients of the decomposition of **1**_*X*_ in the eigenvector basis of **R**_*XX*_, **v**_*i*_ denotes the *i*-th eigenvector of **R**_*XX*_, and *λ*_*i*_ are the corresponding eigenvalues ordered in ascending order.

With the above definitions, the resulting change in the mean activity of cell type *X* induced by the uniform perturbation can then be expressed as:

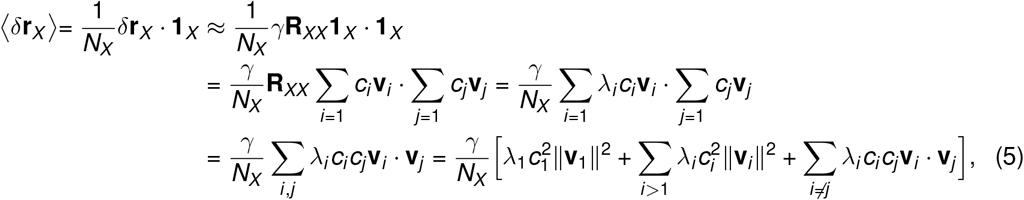

where ⟨·⟩ denotes the average over all neurons of cell type *X*. Even if cell type X is required for stabilization and *λ*_1_ *<* 0, however, due to the heterogeneity of activity in space, 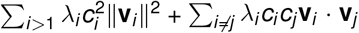 can be larger than 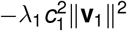 (as shown by the analytical results in Fig. 4A). Therefore, in networks with spatially heterogeneous activity, even if a particular interneuron type is required for stabilization, uniform perturbations of that population do not necessarily induce a paradoxical change in its mean activity.

However, if the real part of the smallest eigenvalue of **R**_*XX*_ is negative (i.e. *Re*(*λ*_1_) *<* 0, Fig. S10), when a patterned perturbation *γ***v**_1_ of cell type *X* is applied in the direction of the eigenvector of **R**_*XX*_ corresponding to that eigenvalue, this patterned perturbation will induce a paradoxical change in the activity of cell type *X* with the change occurring in the direction opposite to the patterned perturbation (Fig. 4B). This behavior can be explained mathematically as follows:

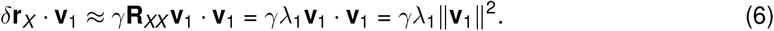

Since λ_1_ is negative, the resulting change in its activity, *δ***r**_*X*_, lies in the direction opposite to the patterned perturbation *γ***v**_1_, indicating a pattern-specific paradoxical effect induced by the patterned perturbation. Note that, in contrast to uniform perturbations, patterned perturbations perturb neurons differently across space. More specifically, patterned perturbations of SST that evoke pattern-specific paradoxical effects perturb SST neurons in the center more strongly than those in the surround (Fig. S11), consistent with a spatially localized contribution of SST neurons to network stabilization (Fig. 3A).

Taken together, although cell-type-specific stabilization cannot be reliably determined from uniform perturbations in our models, it can be revealed through patterned perturbations.

### Recurrent connections and input-output nonlinearities are crucial for context-dependent processing

Next, we aim to determine how recurrent connections shape the integration of feedforward and feedback inputs during context-dependent processing. To this end, we optimize circuit models without recurrent connections. In those optimized models, the size-tuning curves of E and PV populations still exhibit surround suppression (Fig. 5A). However, without recurrent interactions, the stimulus sizes at which the size-tuning curves peak align with those of the L4 and LM inputs (Fig. S12), but differ from the experimental observations (Fig. S1). Furthermore, VIP population displays nearly flat size-tuning curves and spatial response profiles (Fig. 5A), which is inconsistent with experimental data (Fig. S13, Fig. S1). Given that recurrent excitation constitutes the dominant source of excitatory current (Fig. 3, Fig. S5), we then examine the impact of recurrent excitatory connections on spatial responses and size tuning. In the absence of projections from L2/3 excitatory neurons, the E and PV responses to small stimuli fail to reach the experimentally observed levels (Fig. 5B, Fig. S14). This indicates that recurrent excitation is necessary for amplification, particularly when external inputs are weak, as is the case for small stimuli (Fig. S1). Finally, if the rectified quadratic input-output function, which captures the increasing gain with increasing activation observed in cortical neurons that drives the nonlinear response properties of stabilized supralinear networks (Ahmadian et al., 2013; Ahmadian and Miller, 2021), is replaced by a rectified linear function, the top fitted models fail to capture the appropriate E, PV, and VIP size-tuning curves and spatial responses to small stimuli (Fig. 5C, Fig. S15), highlighting the inherently non-linear nature of context-dependent processing. Taken together, these findings highlight the crucial role of recurrent connections and nonlinearities in context-dependent processing.

**Fig. 5.**
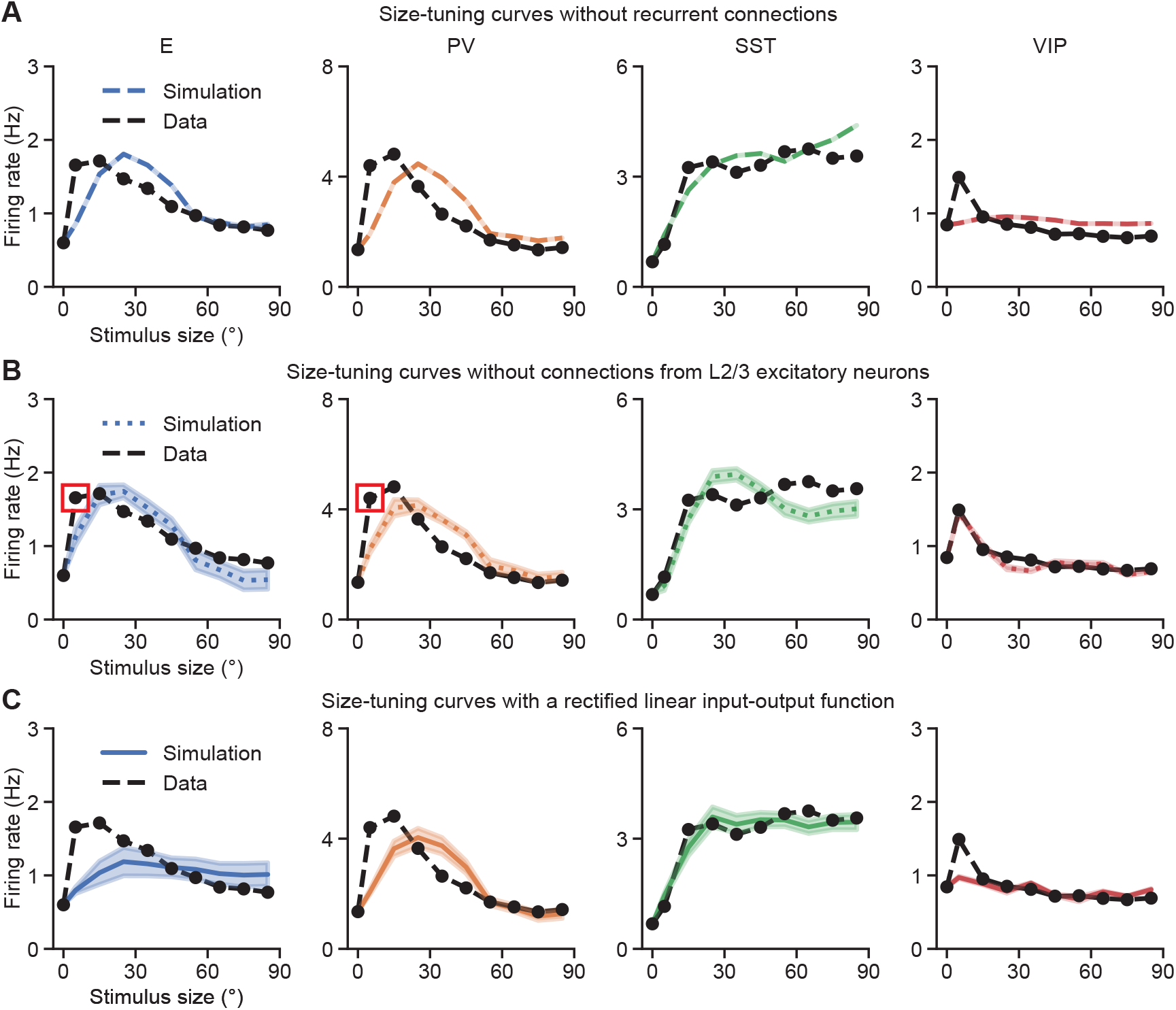
**A**. Size-tuning curves of different cell types from the top ten models obtained by optimizing without recurrent connections. **B**. Same as A, but from the top ten models obtained by optimizing without connections from L2/3 excitatory neurons. The optimized models fail to capture responses to small stimulus sizes, highlighted by the red circles. **C**. Same as A, but from the top ten models obtained by optimization using a rectified linear rather than rectified quadratic inputoutput function.

## Discussion

In this paper, we investigate the circuit characteristics and dynamical signatures that underlie context-dependent processing in spatially extended recurrent neural networks. Using data-driven approaches and analytical methods, we show that the dominance of cell-type-specific inhibition to the excitatory neurons is not static, but dynamically varying with stimulus conditions and spatial location. These changes in the dominance of cell-type-specific inhibition are accompanied by changes in the requirement for SST neurons in network stabilization. Importantly, the resulting changes in cell-type-specific stabilization cannot always be reliably detected using uniform perturbations, instead, they require patterned perturbations that probe the circuit in a spatially dependent manner. Finally, our work highlights the importance of ubiquitous biological features, such as recurrent connections and input-output nonlinearities, in shaping the integration of feedforward and feedback inputs during context-dependent processing.

Due to the limited availability of experimental recordings, we make several assumptions. While inputs from L4 and LM to V1 L2/3 are constrained by the data, we additionally assume that L2/3 receives a residual input with a simple two-dimensional Gaussian spatial profile, whose amplitude increases with stimulus size. Although the current provided by this residual input is relatively weak compared to that from L4 and LM, without the residual input, the optimized models fail to maintain high SST activity at large stimulus sizes (Fig. S7, Fig. S8). As demonstrated in multiple experimental studies, in addition to L4 and LM, V1 L2/3 also receives inputs from other higher visual areas (Froudarakis et al., 2019; Morimoto et al., 2021; Ding et al., 2025) as well as from non-visual brain regions (Zhang et al., 2014; Makino and Komiyama, 2015; Leinweber et al., 2017). Some of these additional inputs may vary in a stimulus-dependent manner (de Vries et al., 2020). To account for these unmeasured contributions, here we introduce the residual input. It will be interesting future work to experimentally identify the joint contributions of brain regions beyond L4 and LM to L2/3.

Our top fitted models naturally capture several experimentally identified features of recurrent circuitry in V1 L2/3 and further reveal additional circuit-level characteristics. First, consistent with experimentally reported connectivity patterns (Pfeffer et al., 2013), the fitted models exhibit neg-ligible connection strengths from SST to SST and from VIP to VIP neurons, and weak strengths from VIP neurons onto E, and PV neurons. Second, the inferred networks operate in an inhibition-stabilized regime already at baseline, even in the absence of sensory drive (i.e., stimulus size 0°, Fig. 3C). This behavior aligns with experimental evidence indicating that cortical circuits are inhibition-stabilized during spontaneous activity (Sanzeni et al., 2020). Our models further demonstrate that PV neurons are required to stabilize the network across all stimulus sizes, whereas SST-mediated stabilization is also required, likely only for larger sizes but with a modest degree of heterogeneity across models (Fig. S9). Whereas recent studies have implicated SST-mediated stabilization in response to increasing stimulus contrast during locomotion (Waitzmann et al., 2024; Cammarata et al., 2025), our work shows that in spatially extended networks, SST-mediated stabilization is required as stimulus size increases while contrast remains unchanged. This may seem to be contradicted by recent experiments that found that suppressing PV activity led to runaway cortical activity, but suppressing SST activity did not, even at the largest stimulus sizes (Veit et al., 2017). However, the same is true in our model: in contrast to silencing PV, silencing SST neurons for large stimulus sizes mainly results in elevated but still physiologically plausible activity levels (Fig. S16). This indicates that SST as well as PV are stabilizing the response to the visual stimulus, but without SST the network moves to a different, solely PV-stabilized visual response with higher firing rates. This suggests that PV neurons play the dominant role in stabilizing the network.

Experimentally, inhibition stabilization is commonly assessed by examining paradoxical effects using optogenetic perturbations. These perturbations are typically applied uniformly and non-specifically, with all targeted neurons receiving the same perturbative input. Under such uniform perturbations, paradoxical effects are defined as a decrease (increase) in the mean activity of inhibitory neurons in response to direct excitatory (inhibitory) input to them. Consistent with previous experimental findings (Sanzeni et al., 2020), uniform perturbations of the PV population in our fitted spatially extended models robustly elicit paradoxical responses within the PV population. In contrast, applying the same uniform perturbations to the SST population does not reliably induce paradoxical effects, even when SST neurons are required for network stabilization. Our work suggests that a more reliable assessment of cell-type-specific stabilization requires patterned perturbations that are tailored to the spatial structure of network activity. Importantly, the patterned perturbations predicted by our model exhibit a specific spatial profile and preferentially target SST neurons in the center more strongly than those in the surround (Fig. S11), consistent with a spatially localized role of SST neurons in stabilizing network dynamics. Patterned perturbations induce pattern-specific paradoxical effects, in which the activity of the perturbed cells changes in the direction opposite to the applied perturbation. These predictions could be directly tested in future experiments using holographic optogenetic perturbations.

Patterned perturbations have been previously proposed as an approach to revealing functionally specific inhibition-stabilized networks, in which specific unstable excitatory subnetworks are sta-bilized by dedicated inhibitory subnetworks (Sadeh and Clopath, 2020). Here, in networks with multiple interneuron subtypes and spatially heterogeneous activity, we show that patterned perturbations unmask paradoxical responses that become undetectable under uniform perturbations, and can therefore more robustly reveal cell-type-specific stabilization properties. Note that while the relationship between inhibition stabilization and paradoxical effects has been extensively investigated in low-dimensional population models (Tsodyks et al., 1997; Sanzeni et al., 2020; Mahrach et al., 2020; Wu and Gjorgjieva, 2023; Waitzmann et al., 2024), this relationship is more nuanced in high-dimensional networks (Miller and Palmigiano, 2020; Shao et al., 2025). As demonstrated by (Miller and Palmigiano, 2020), in high-dimensional networks, the number of unstable modes in the subnetwork excluding a given cell type should have the same parity as the number of patterned perturbations of that cell type that elicit paradoxical responses (paradoxically responding modes) in the excluded population (see Methods). Consistent with this, our models exhibit the same parity relationship, despite these two numbers not necessarily being identical (Fig. S17, Fig. S18).

Our models show that recurrent excitation provides the dominant source of excitation, implying a potentially important role of recurrence in context-dependent processing. Indeed, this is directly supported by the fact that circuit models optimized without recurrent connections, and in particular without connections from L2/3 excitatory neurons, fail to produce sufficient response amplitudes for small stimulus sizes. This highlights the crucial role of recurrent excitation in response amplification, consistent with recent experimental findings (Peron et al., 2020; Jennings et al., 2025). Furthermore, the biologically realistic expansive input-output nonlinearities are critical for producing sufficient response amplitudes for a wider range of smaller stimulus sizes, suggesting the importance of this nonlinearity for response amplification.

In summary, our work reveals stimulus-dependent stabilization and perturbation response signatures of recurrent cortical circuits with diverse cell types, and highlight the importance of ubiquitous biological components such as recurrent connections and expansive input-output nonlinearities in context-dependent processing.

## Methods

### Data

We used data from (Keller et al., 2020a,b) measuring activity from V1 L4 excitatory, V1 L2/3 excitatory, PV, SST, and VIP neurons, as well as LM boutons in V1 L1, using two-photon calcium imaging in awake, head-fixed mice viewing drifting grating stimuli at 100% contrast. In total, recordings included 1,489 L2/3 excitatory neurons, 25 PV neurons, 105 SST neurons, 90 VIP neurons, 38 L4 excitatory neurons, and 167 LM excitatory boutons. Stimuli were presented with sizes ranging from 5°to 85°, in 10°increments. The authors determined the distance between each neuron’s receptive field center and the stimulus center from receptive field mapping experiments. Further experimental details are available in (Keller et al., 2020a,b). We used the analysis of responses from (Di Santo et al., 2025), which defined response amplitude as the mean response over the 2-s stimulus presentation period, converted from fluorescence signals to firing rates using the procedure described in (Dipoppa et al., 2018), and constructed rate fields (mean rate vs. spatial position) for each cell type.

### Model

To optimize the spatially extended circuit model to reproduce the experimentally observed responses, we discretize the model on a regular two-dimensional grid with 30 × 30 spatial locations along the horizontal and vertical dimensions, respectively. The grid spacing is 6°along both spatial dimensions, such that each unit represents the average activity within a 6°×6°region of the retinotopic coordinate space. The rate dynamics of a unit of cell type *A* at grid location **x** in response to a stimulus of size *s* is given by:

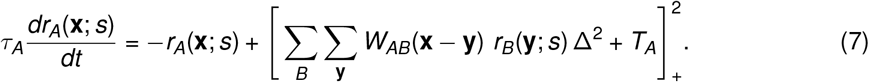

Here, *τ*_*A*_ is the time constant of the rate dynamics. *W*_*AB*_(**x**−**y**) denotes the connection strength from a unit of cell type *B* at grid location **y** to a unit of cell type *A* at grid location **x**. The factor Δ^2^ corresponds to the area of a single grid element and arises from the discretization of the continuous spatial integral. *T*_*A*_ is the bias, and 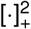 denotes a rectified quadratic function. The stimulus size, *s*, is varied from 5°to 85°in increments of 10°. Cell types are defined as *A* ∈ {*E, PV, SST, VIP*} and *B* ∈ {*E, PV, SST, VIP, L*4, *LM, R*}. Here, *R* is the residual input, representing input from unmeasured sources. The feedforward input from L4 and the feedback input from LM are determined by experimental data, while the residual input is modelled by a simple 2D Gaussian with fixed spatial scale and a peak amplitude that increases with the square root of the stimulus size *s* (A linear dependence on *s* would lead to a 17-fold amplitude increase for large vs. small stimuli; the square root scaling was chosen to reduce this):

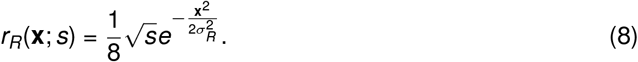

The spatial profiles of *W* are modelled by 2D isotropic Gaussians,

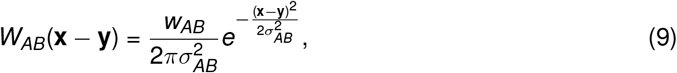

with the spatial scale *σ* extracted from previous literature (Table 1; for detailed descriptions, please see Di Santo et al., 2025).

The connection strengths *w*_*AB*_ and biases *T* are optimized using backpropagation through time to minimize the error between simulated and experimentally measured responses of the recurrent cell types. Following Chau et al. (2025), the optimization is performed using the scipy.optimize.minimize function with the SLSQP (Sequential Least Squares Programming) algorithm, with gradients computed using PyTorch’s automatic differentiation engine. Specifically, the loss function *L* subject to optimization is defined below:

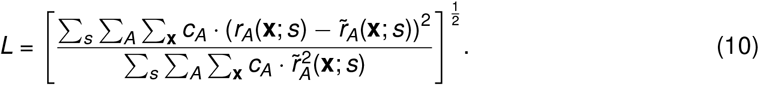

Here, *r*_*A*_(**x**; *s*) is the simulated response of a unit located at grid location **x** of cell type *A* to a stimulus of size *s* in the model. The location **x** specifies the unit’s receptive field center, allowing the distance between its receptive field center and the stimulus center in retinotopic space to be computed. 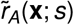 is the corresponding response observed in the data. The sum is over *A* ∈ {*E, PV, SST, VIP*}, and *c*_*A*_ is a non-negative weighting factor assigned to cell type *A* (given in Table 1), allowing greater weight to be given to cell types found to be more difficult to fit. Note that the denominator serves merely as a normalization factor, for the more general purpose beyond our current work of allowing comparison of losses on different data sets, and does not affect the optimization.

### Cell-type-specific stabilization

To access cell-type-specific stabilization properties of the recurrent network, we express the dynamics of the recurrent network in the following general form (Miller and Palmigiano, 2020):

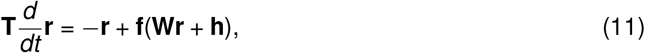

where **T** is a diagonal matrix containing the time constants of rate dynamics for neurons across different recurrent cell types, **r** is a vector containing the firing rates, **f** is a rectified quadratic function, applied elementwise. **W** is the connectivity matrix, **h** is the effective input vector, equivalent to the sum of currents from external inputs and bias terms.

The Jacobian of the recurrent network is given by:

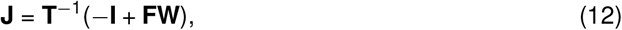

where **I** is the identitiy matrix, and **F** is a diagonal matrix containing the derivatives of corresponding input-output functions evaluated at the fixed point. We break the matrices **T, F, I**, and **W** into their cell-type-specific submatrices, **T**_*A*_, **F**_*A*_, **I**_*A*_, and **W**_*AB*_, with *A, B* ∈ {*E, P, S, V*}. Note that the nonzero entries of the diagonal matrices **T**_*A*_, **F**_*A*_, and **I**_*A*_ are positive, while the entries of *W*_*AB*_ are non-negative for *B* = *E* and non-positive for *B* ∈ {*P, S, V*}. Then the Jacobian can be written:

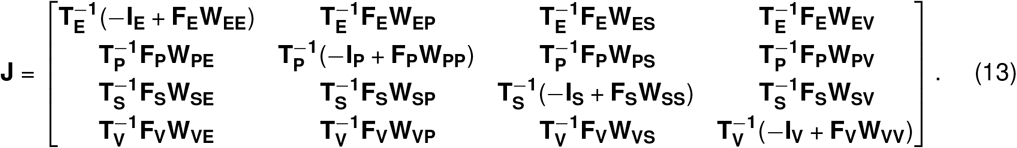

To assess whether specific inhibitory populations are necessary for stabilization, we compute the Jacobian of subnetworks in which selected inhibitory populations are excluded. If the leading eigenvalue of this Jacobian has a positive real part, this indicates that the excluded population is required for stabilization. For every stimulus size, we evaluate the Jacobian of four different subnetworks at the steady state: of the E subnetwork to assess whether inhibition in general is required for stabilization; and of the network with PV, SST, or VIP populations excluded, to determine if each is required for stabilization. The Jacobian of the E subnetwork is given by the top left entry of Eq. 13, while the Jacobian of the subnetwork with one cell type excluded is given by Eq. 13 with the column and row corresponding to the given cell type removed.

### Paradoxical effects

Assume that for input **h**^*\**^, the network has a fixed point **r**^*\**^:

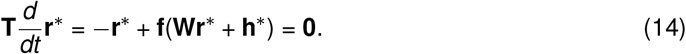

Now consider a small perturbation *δ***h** to the input, after which the network converges to a new fixed point **r**^*\**^ + *δ***r**,

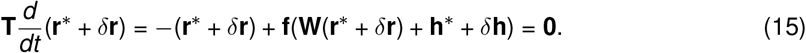

Taking a first-order Taylor expansion of **f**(**W**(**r**^*\**^ + *δ***r**) + **h**^*\**^ + *δ***h**) about **Wr**^*\**^ + **h**^*\**^, we can express the resulting change in network activity, *δ***r**, as a function of the input perturbation, *δ***h**, as:

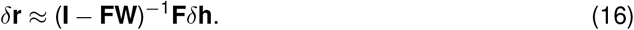

We further define the response matrix **R** as follows:

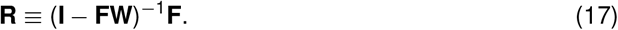

Similarly, the change in activity of cell type *X, δ***r**_*X*_, induced by a change in their input, *δ***h**_*X*_, can now be written as:

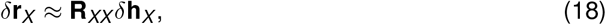

where **R**_*XX*_ denotes the *XX* submatrix of **R**.

A paradoxical response to the perturbation is one for which

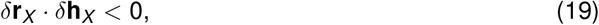

where · represents the dot product of two vectors. Note that this condition indicates that the change in activity of cell type *X* has a component pointing opposite to the applied perturbation.

A non-paradoxical response is one for which

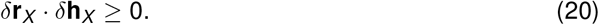

To demonstrate the effects of uniform and patterned perturbations on neural activity, we denote **V** as the matrix whose columns are the eigenvectors of **R**_*XX*_, and Λ as the diagonal matrix of corresponding eigenvalues:

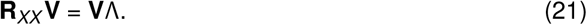

For the *i*-th eigenvector **v**_*i*_ and *i*-th eigenvalue *λ*_*i*_, we then have

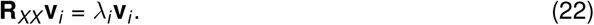

A uniform excitatory perturbation to cell type *X* can be written as *γ***1**_*X*_, where *γ >* 0 controls the perturbation amplitude and **1**_*X*_ denotes the all-ones vector. The **1**_*X*_ can be expressed as a linear combination of the basis vectors in **V**:

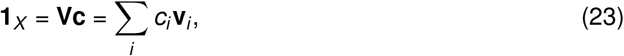

where the coefficient vector, **c**, can be obtained by

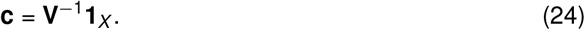

The resulting change in the activity of cell type *X* induced by the uniform perturbation can therefore be written as:

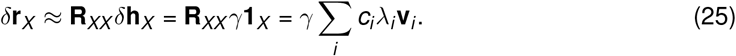

A patterned perturbation to cell type *X* that is in the same direction as the eigenvector corresponding to the smallest eigenvalue of **R**_*XX*_ can be written as *γ***v**_1_, where *γ >* 0 controls the perturbation amplitude. The resulting change in the activity of cell type *X* induced by this patterned perturbation can therefore be written as:

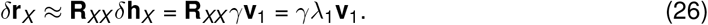

### Relationship between cell-type-specific stabilization and paradoxical effects

A fixed point is stable if and only if all eigenvalues of the Jacobian **J** computed at the fixed point have negative real parts. Consequently, all eigenvalues of −**J** have positive real parts, and so det(−**J**) *>* 0. We denote the Jacobian of the subnetwork excluding cell type *X* by **J**_−*X*_. As shown in (Miller and Palmigiano, 2020), det(−**J**_−*X*_) and det(**R**_*XX*_) have the same sign, which implies that an odd (even) number of unstable modes in the subnetwork excluding cell type *X* (eigenvectors of −**J**_−*X*_ corresponding to eigenvalues with negative real parts) corresponds to an odd (even) number of modes of cell type *X* that respond paradoxically (eigenvectors of **R**_*XX*_ corresponding to eigenvalues with negative real parts). Consistent with this, our results show that the parity of the number of unstable modes in the subnetwork excluding cell type *X* matches the parity of the number of paradoxically responding modes of cell type *X* (Fig. S17, Fig. S18). Importantly, despite having the same parity, these numbers can have different values (Fig. S17B).

## Data Availability

The code used for model simulations is available at https://github.com/yuekriswu/context-dependentprocessing.

## Contributions

Conceptualization, K.D.M., Y.K.W., and S.D.S.; methodology, Y.K.W.; software, Y.K.W. and H.Y.C.; mathematical analysis, Y.K.W. and K.D.M.; writing-original draft, Y.K.W.; writing-review & editing, K.D.M. and Y.K.W.; visualization, Y.K.W.; supervision, K.D.M.; funding acquisition, K.D.M.

## Acknowledgments

We thank Andreas Keller, Morgane Roth, and Massimo Scanziani for generously providing their data. This work is supported by the Gatsby Charitable Foundation (GAT3708), the NIH (1RF1DA056397, U19NS107613, and T32 EY013933 to H.Y.C.), the NSF (DGE-2036197 to H.Y.C.), and the Simons

Foundation (SCGB 543017 to KDM). S.D.S. was supported by the Agencia Estatal de Investigación (AEI) through the Project of I+D+i (Ref. PID2023-149174NB-I00), financed by MICIN/AEI/10.13039/501100011033 and the ERDF/EU.

## Supplementary Figures

**Fig. S1.**
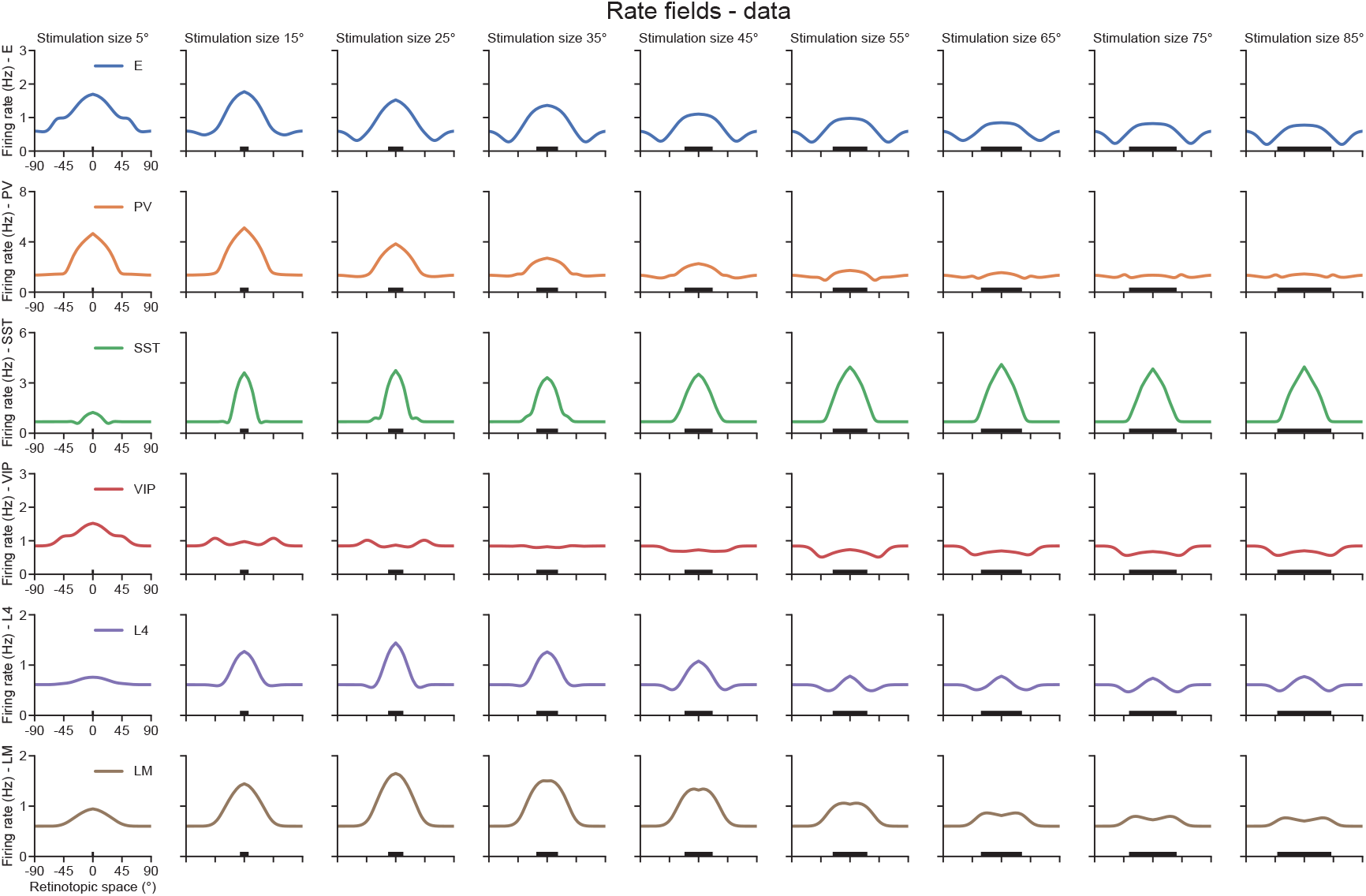
Rate field data of E, PV, SST, VIP, L4, and LM populations across stimulus sizes ranging from 5°to 85°. Black bars indicate stimulus size. Adapted from Di Santo et al., 2025.

**Fig. S2.**
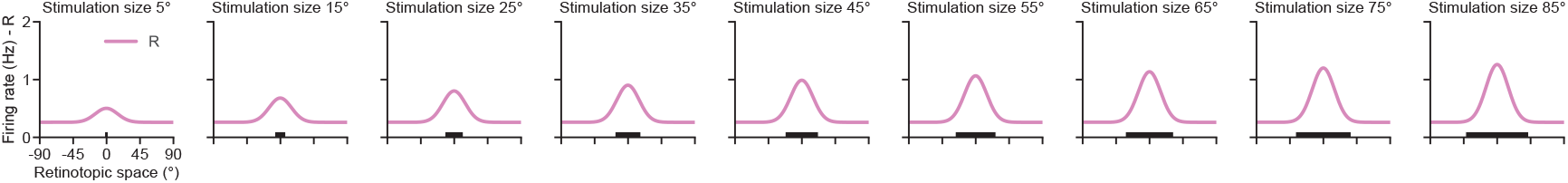
Constructed rate fields of the residual input across stimulus sizes. The peak amplitude of the rate field increases with stimulus size, while its spatial scale *σ*_*R*_ (see Eq. 8) remains fixed at 15°across all stimulus conditions.

**Fig. S3.**
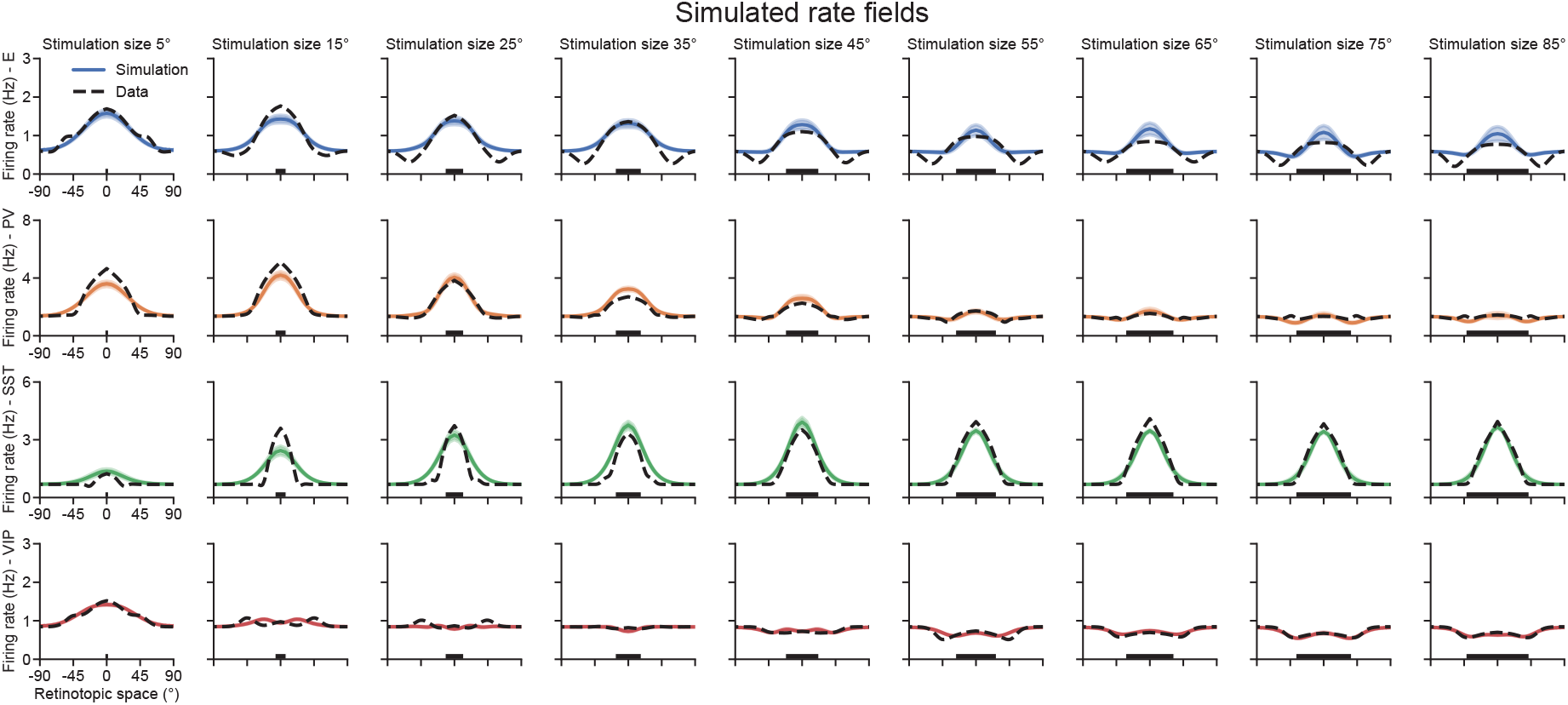
Simulated rate fields of E, PV, SST, VIP populations across stimulus sizes from the top ten optimized models. Solid lines and shaded regions indicate the mean and the standard deviation of the simulated rate fields, respectively. Dashed lines represent the corresponding data.

**Fig. S4.**
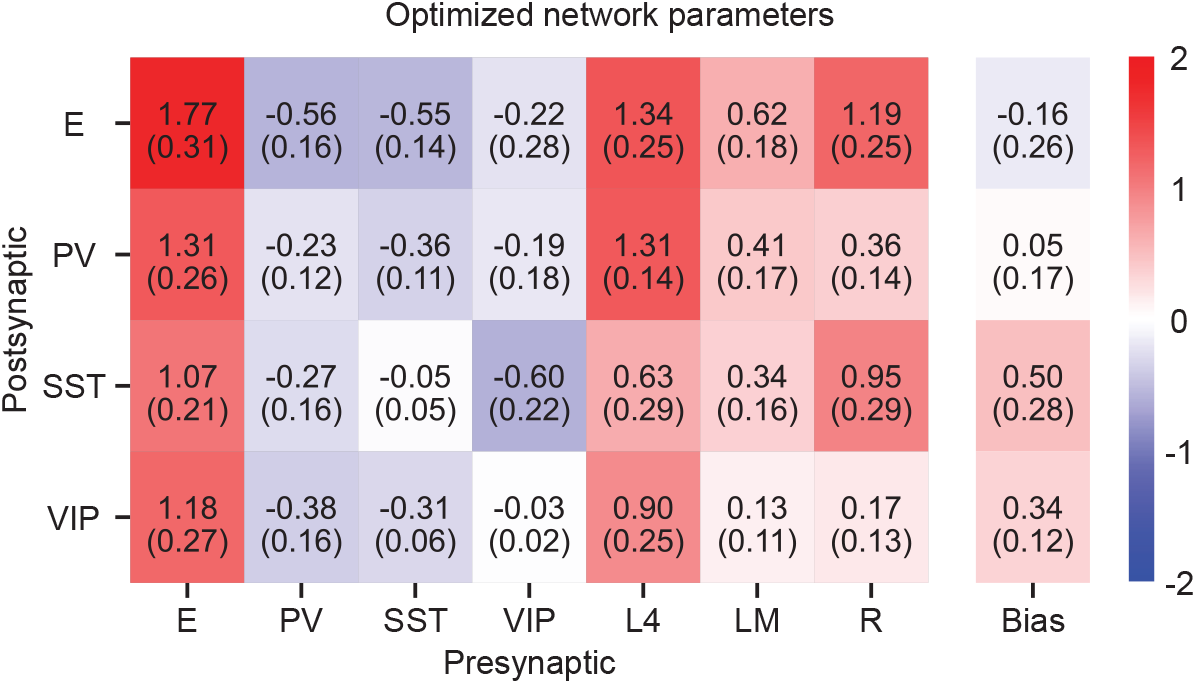
Connection strengths and biases from the top ten optimized models. The mean values and standard deviations are shown above and below, respectively. Color codes the mean value.

**Fig. S5.**
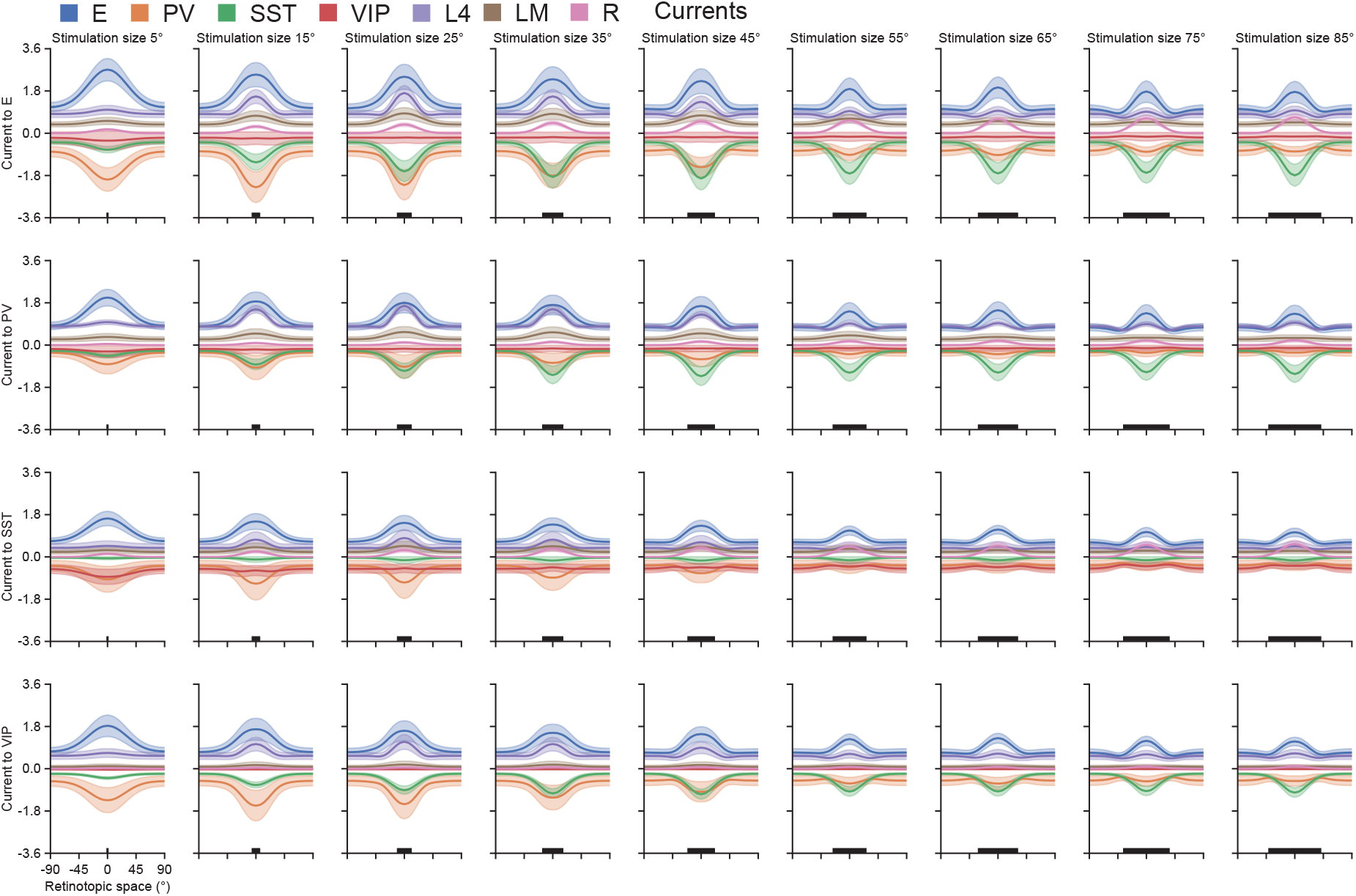
Different current sources to E, PV, SST, and VIP populations across stimulus sizes. Solid lines and shaded regions indicate the mean and the standard deviation, respectively, of the corresponding currents from the top ten optimized models.

**Fig. S6.**
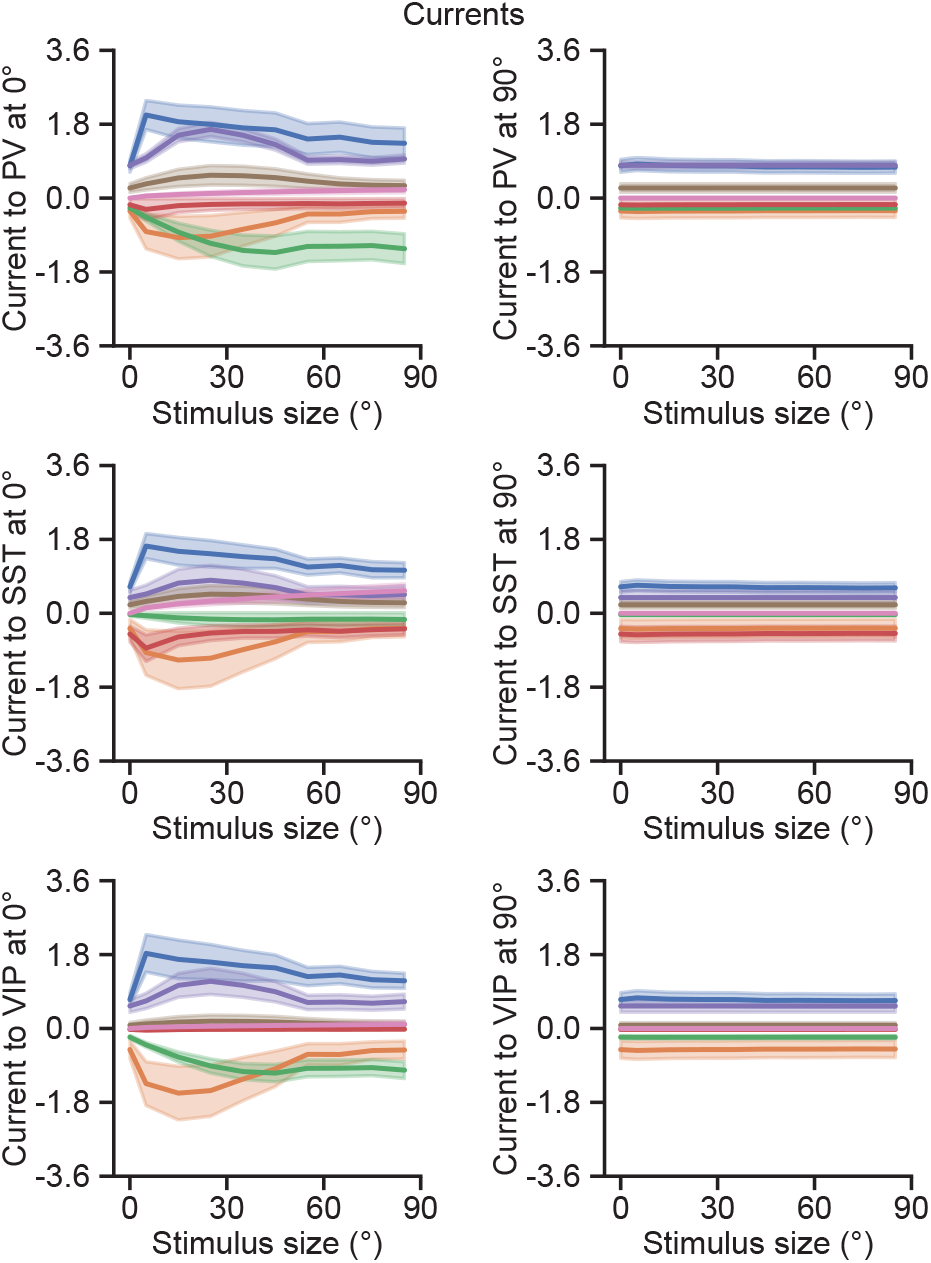
Left: Current from different sources to different types of inhibitory neurons (top: PV, middle: SST, bottom: VIP) located at 0° in the retinotopic space as a function of stimulus size. Solid lines and shaded regions indicate the mean and the standard deviation of the corresponding currents, respectively. Right: Same as left but for different types of inhibitory neurons located at 90° in the retinotopic space.

**Fig. S7.**
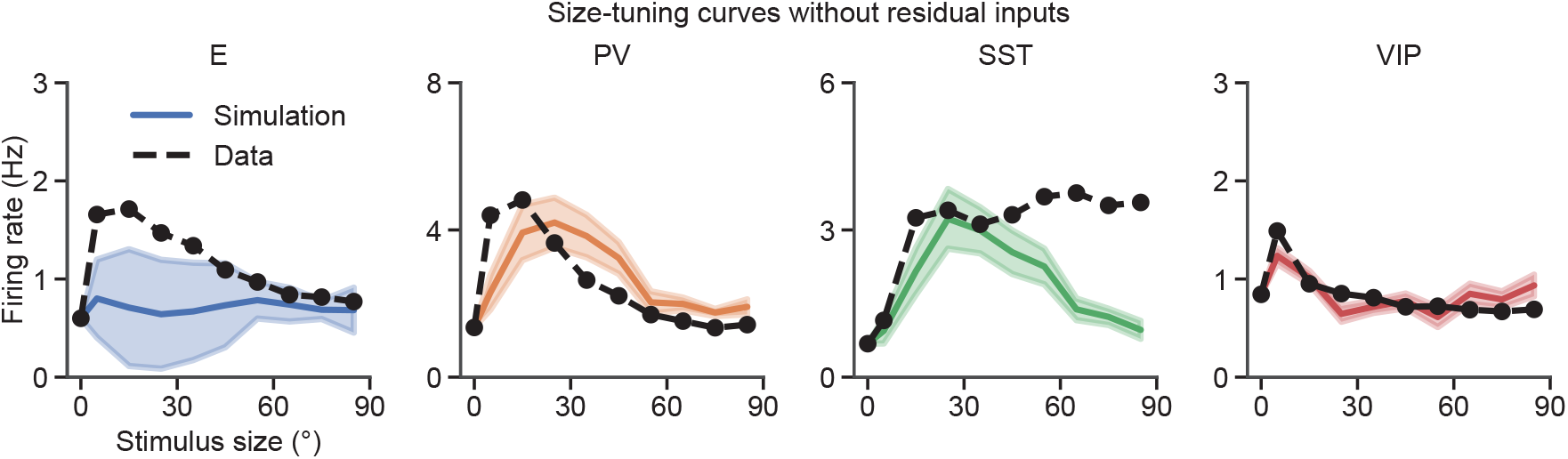
Size-tuning curves of E, PV, SST, and VIP populations from the top ten models obtained by optimizing without residual inputs. Solid lines and shaded regions indicate the mean and the standard deviation of the size-tuning curves, respectively. Dashed lines represent the corresponding data.

**Fig. S8.**
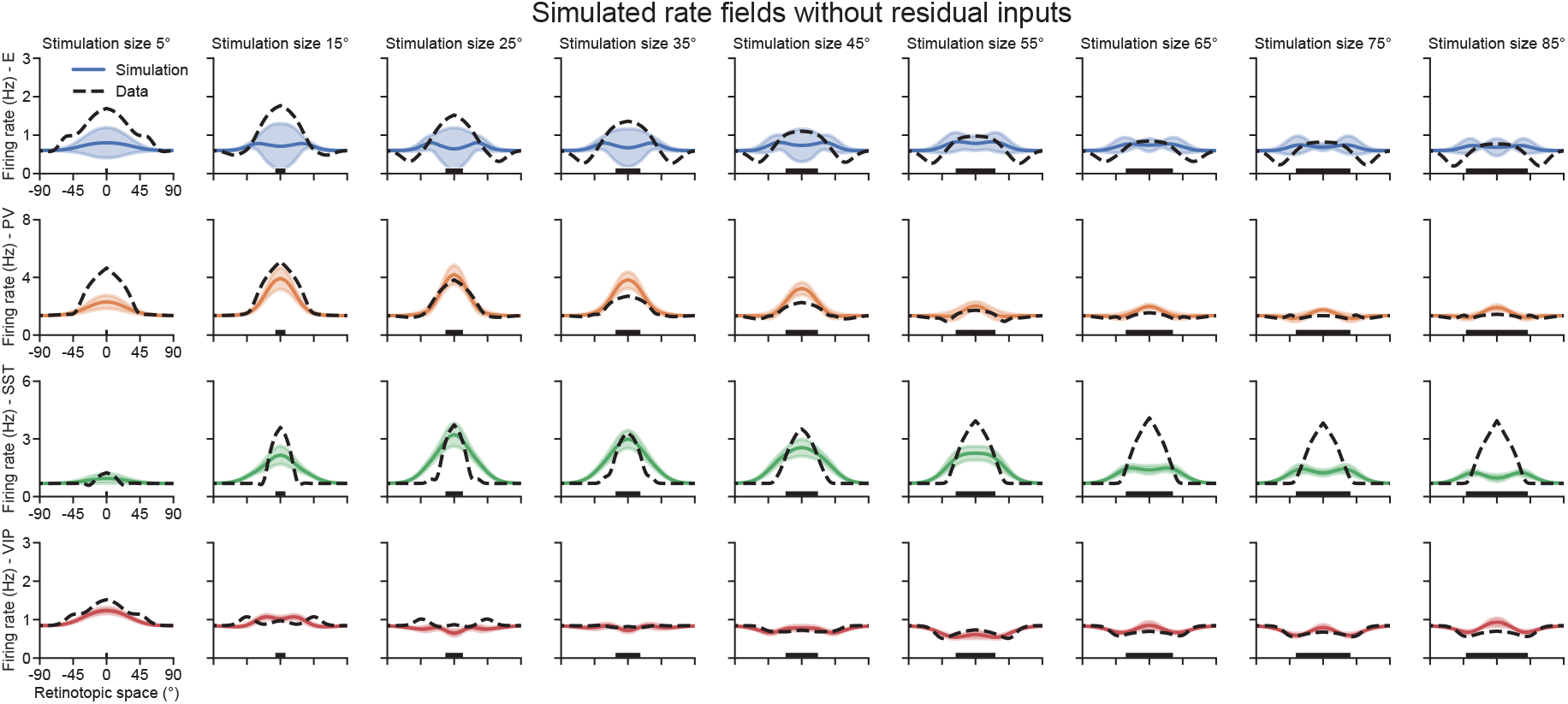
Simulated rate fields of E, PV, SST, VIP populations across stimulus sizes from the top ten models obtained by optimizing without residual inputs. Solid lines and shaded regions indicate the mean and the standard deviation of the simulated rate fields, respectively. Dashed lines represent the corresponding data.

**Fig. S9.**
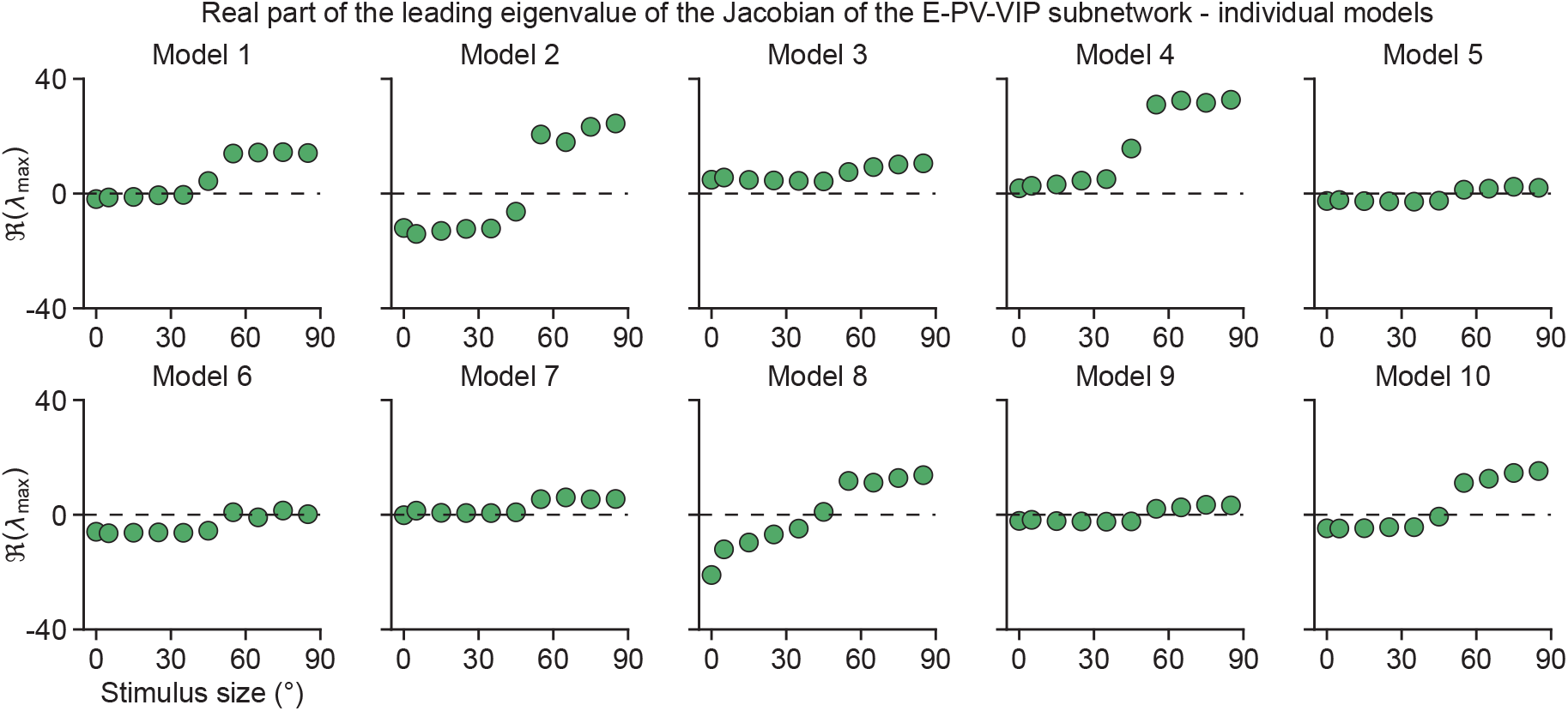
Real part of the leading eigenvalue of the Jacobian of the E-PV-VIP subnetwork as a function of stimulus size. Each subplot corresponds to one model.

**Fig. S10.**
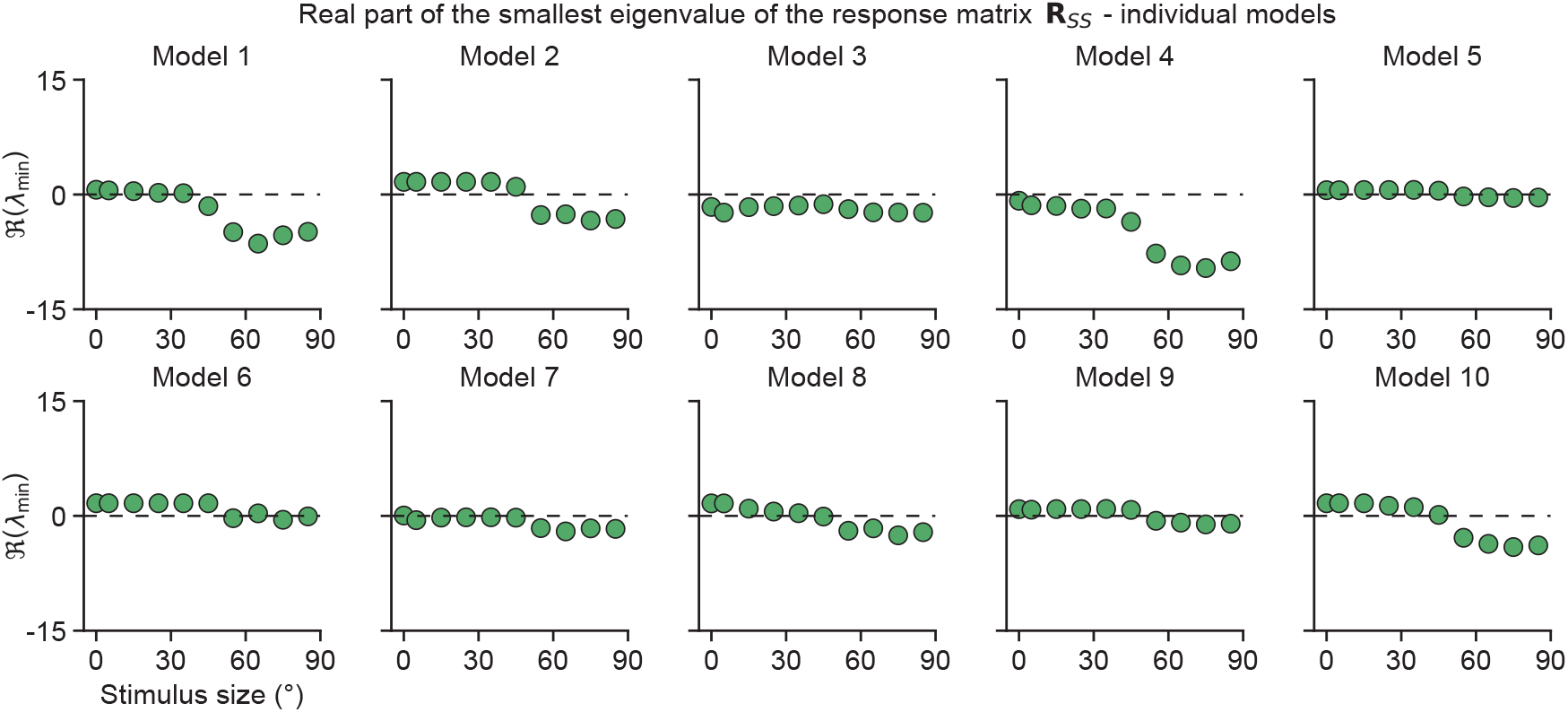
Real part of the smallest eigenvalue of the response matrix **R**_*SS*_ as a function of stimulus size. Each subplot corresponds to one model.

**Fig. S11.**
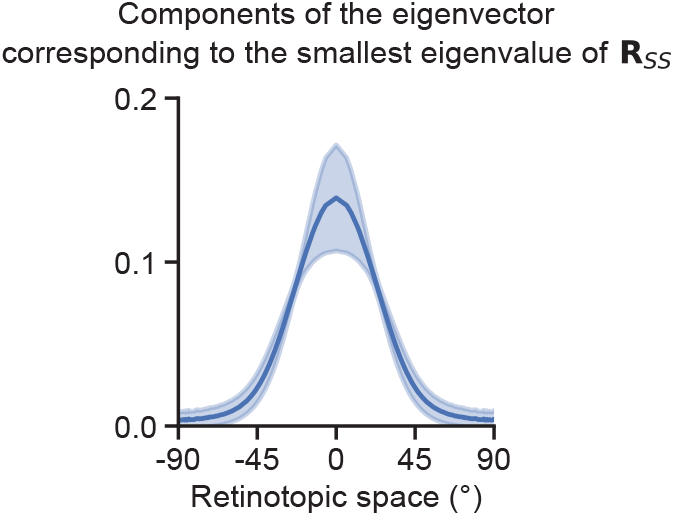
Components of the eigenvector corresponding to the smallest eigenvalue of the response matrix **R**_*SS*_ as a function of retinotopic space. Solid lines and shaded regions indicate the corresponding mean and the standard deviation. Eigenvectors are computed at a stimulus size of 55°, where the real parts of the smallest eigenvalues of the response matrix **R**_*SS*_ are negative for all ten models (Fig. S10).

**Fig. S12.**
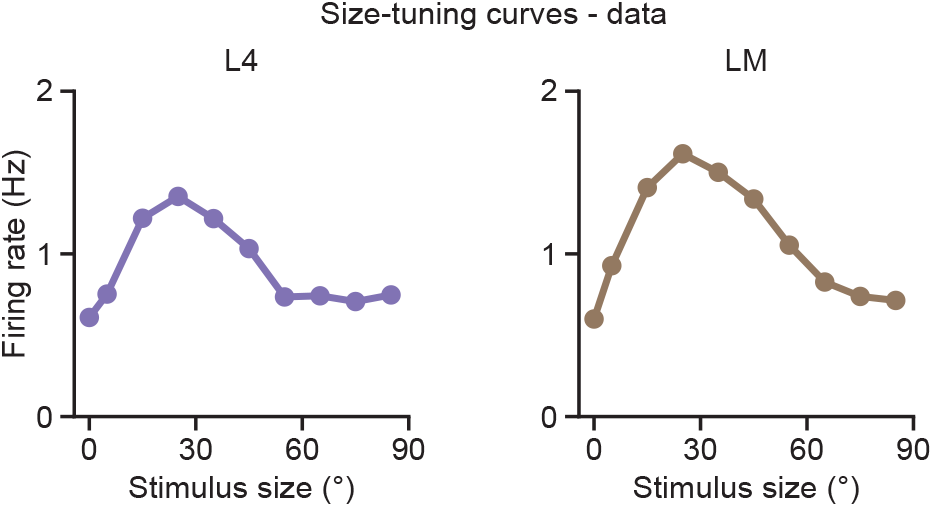
Size-tuning curve data of L4 (left) and LM (right).

**Fig. S13.**
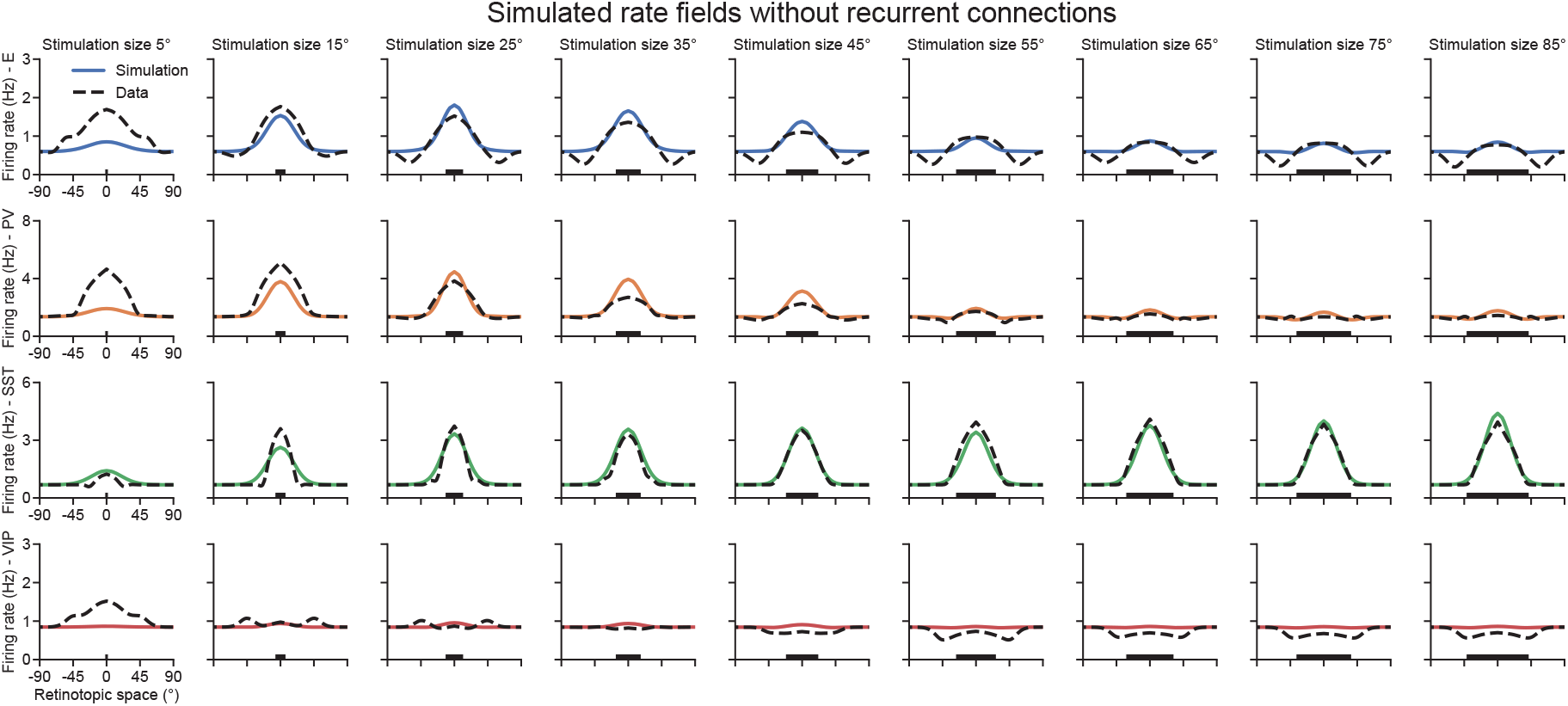
Simulated rate fields of E, PV, SST, and VIP populations across stimulus sizes from the top ten models obtained by optimizing without recurrent connections.

**Fig. S14.**
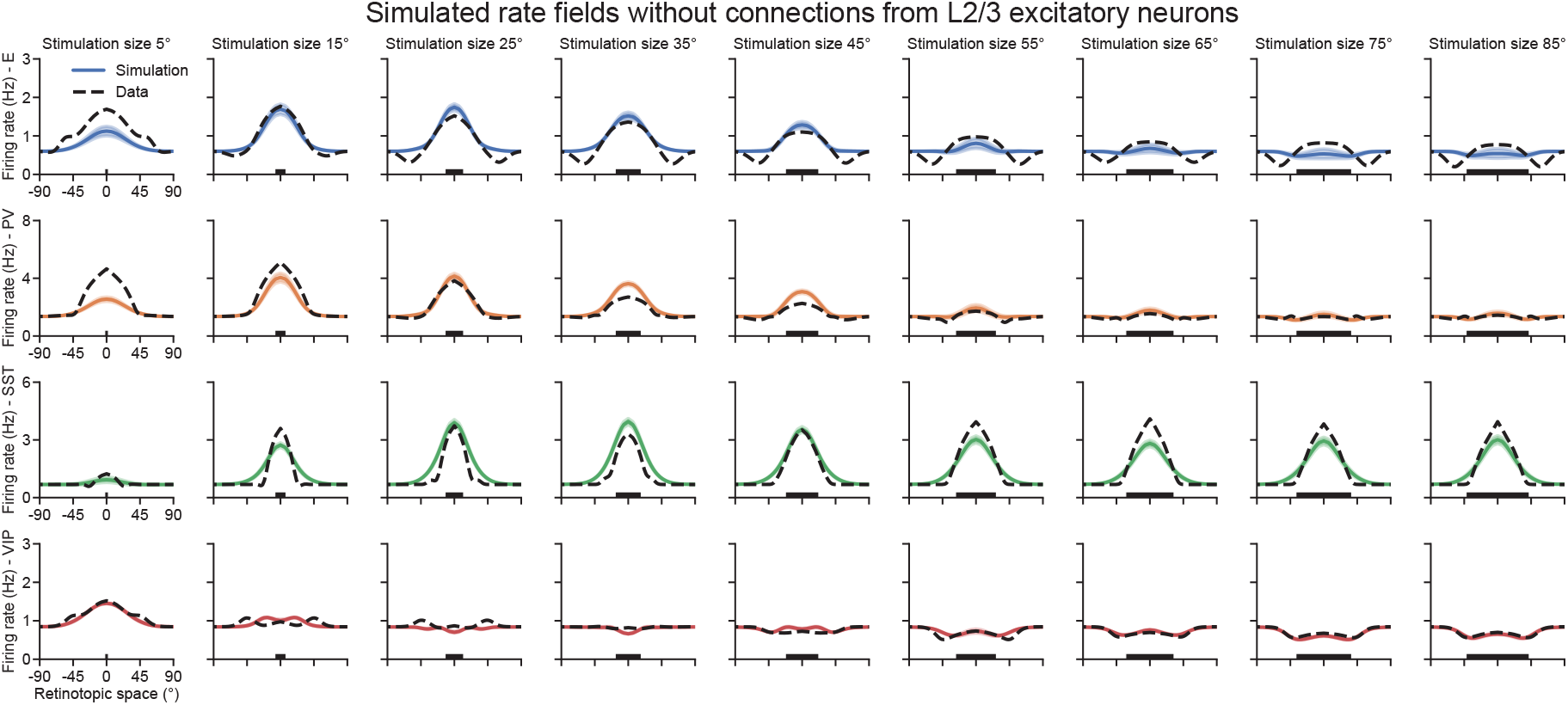
Simulated rate fields of E, PV, SST, and VIP populations across stimulus sizes from the top ten models obtained by optimizing without connections from L2/3 excitatory neurons. Solid lines and shaded regions indicate the mean and the standard deviation of the simulated rate fields, respectively. Dashed lines represent the corresponding data.

**Fig. S15.**
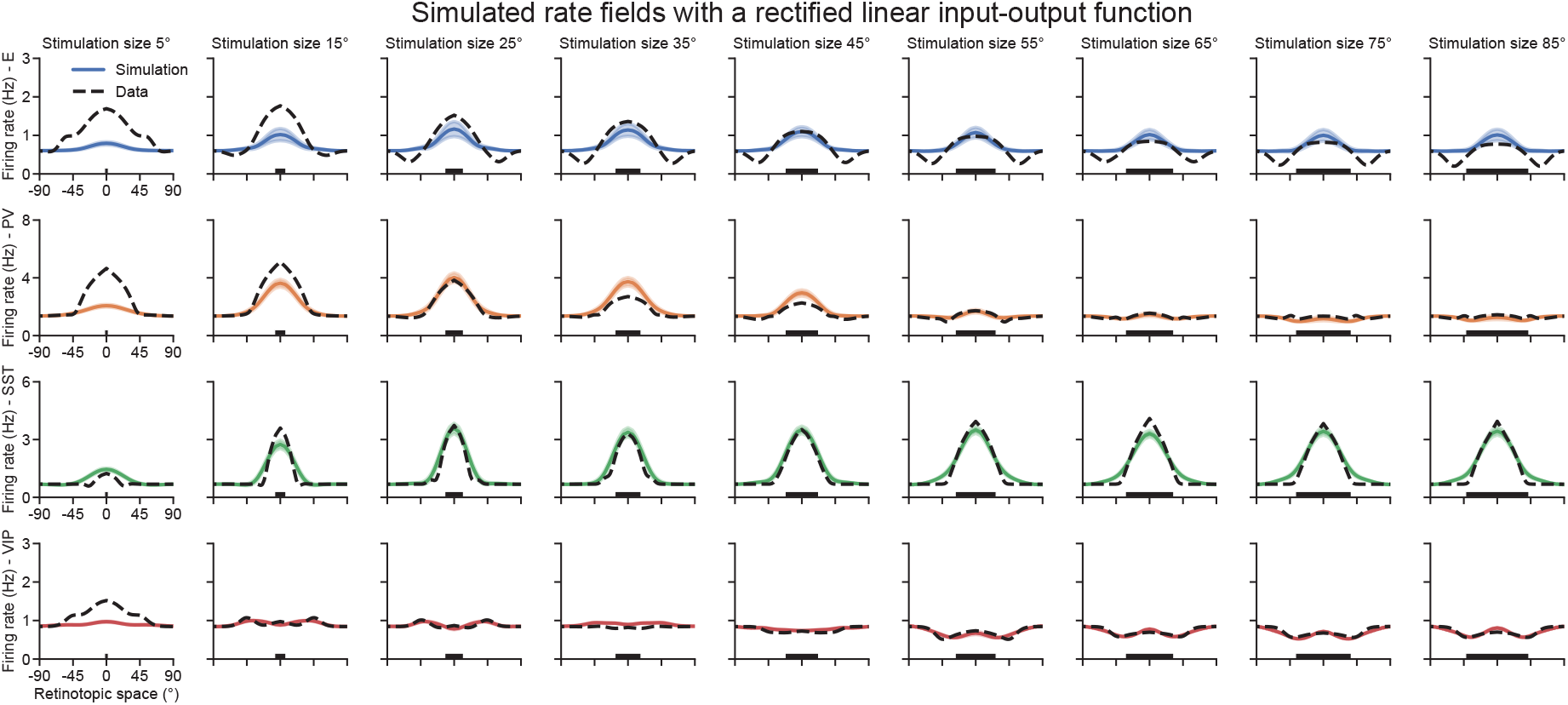
Simulated rate fields of E, PV, SST, and VIP populations across stimulus sizes from the top ten models obtained by optimizing using a rectified linear rather than rectified quadratic input-output function. Solid lines and shaded regions indicate the mean and the standard deviation of the simulated rate fields, respectively. Dashed lines represent the corresponding data.

**Fig. S16.**
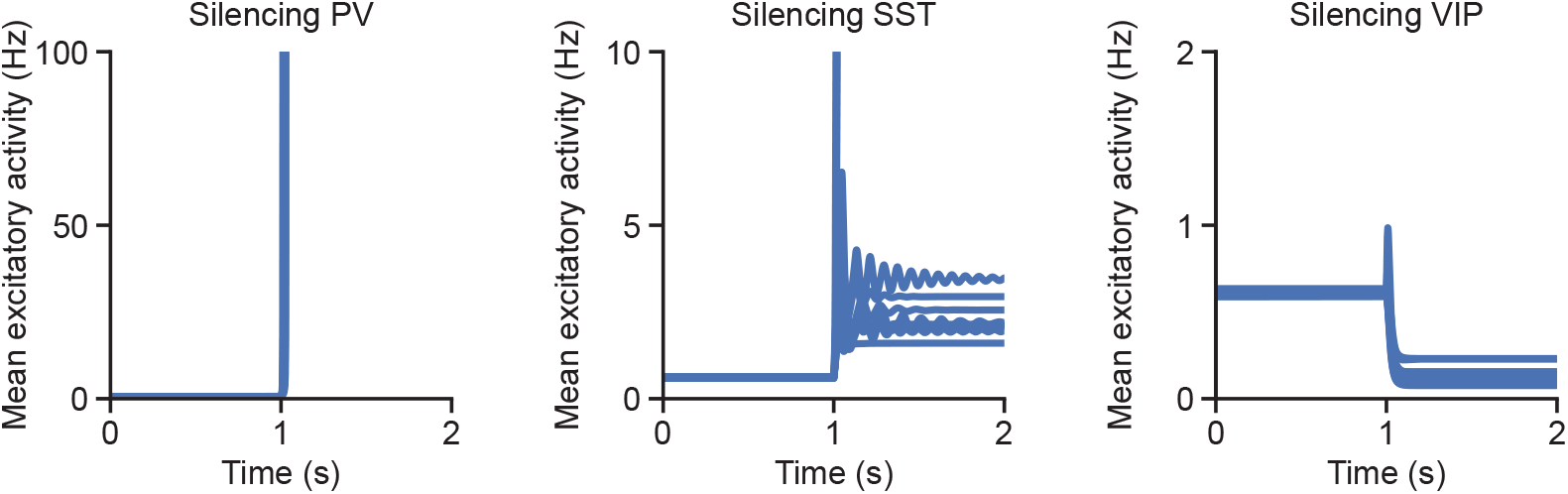
Left: Mean excitatory activity following PV silencing at 1 s for stimulus size 55°. Each line corresponds to one model. In all ten models, excitatory activity exhibits runaway dynamics after PV silencing. Middle: Same as left, but with SST silencing. In eight out of ten models, excitatory activity remains within physiologically realistic levels after SST silencing. Right: Same as left, but with VIP silencing.

**Fig. S17.**
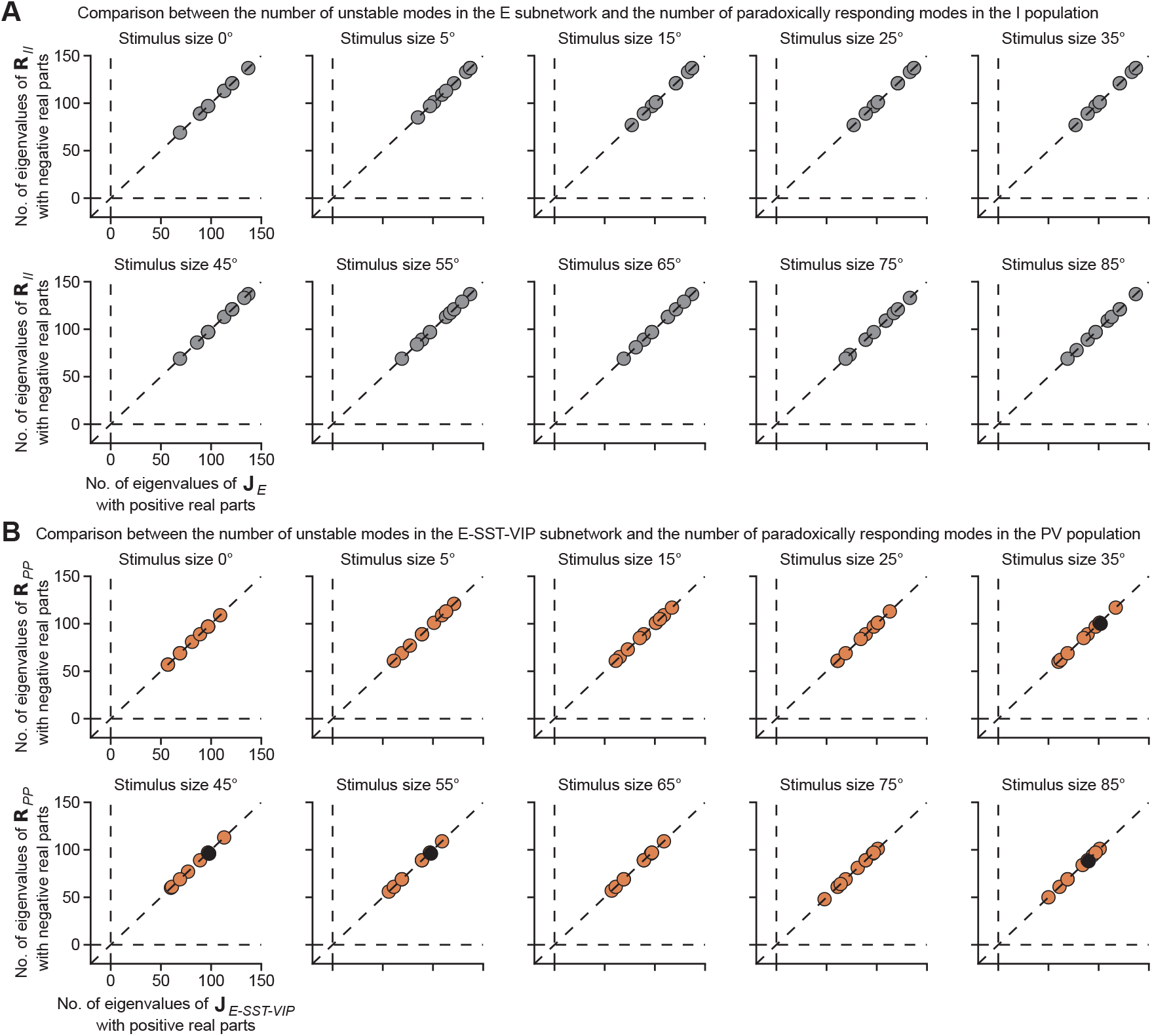
**A**. Comparison between the number of unstable modes in the E subnetwork and the number of paradoxically responding modes in the I population. The number of unstable modes in the E subnetwork is quantified by counting the eigenvalues of the Jacobian of the E subnetwork **J**_*E*_ with positive real parts shown on the x-axis. The number of paradoxically responding modes in the I population is quantified by counting the eigenvalues of the corresponding response matrix **R**_*II*_ with negative real parts shown on the y-axis. Results for different stimulus sizes are shown in separate panels. Each dot represents one of the ten models. All dots have equal, positive x and y values and lie along the diagonal lines, indicating that the parity of the number of unstable modes in the E subnetwork matches the parity of the number of paradoxically responding modes in the I population. Note that some dots are located at the same coordinates. **B**. Same as A, but for the comparison between the number of unstable models in the E-SST-VIP network and the number of paradoxically responding modes in the PV population. The numbers are quantified by the eigenvalues of the Jacobian of the E-SST-VIP subnetwork **J**_*E*−*SST* −*VIP*_ with positive real parts shown on the x-axis and the eigenvalues of the corresponding response matrix **R**_*PP*_ with negative real parts shown on the y-axis, respectively. Some dots do not lie exactly on the diagonal and are colored black (corresponding to x-y pairs of 102-100 at a stimulus size of 35°, two points at 98-96 for 45°, one point at 98-96 for 55°, and one point at 90-88 for 85°), indicating that despite having the same parity, these two values need not be identical.

**Fig. S18.**
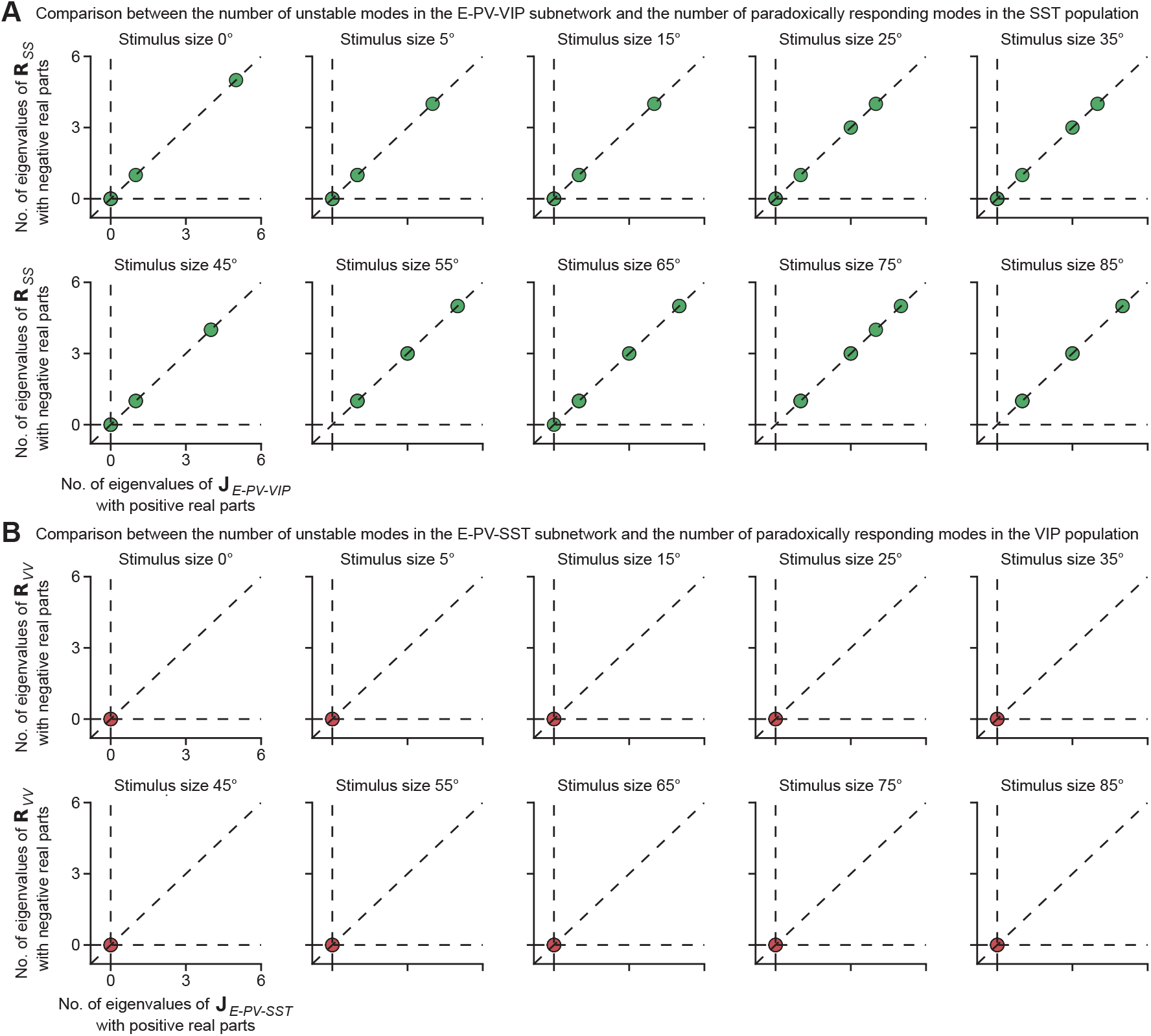
**A**. Comparison between the number of unstable models in the E-PV-VIP subnetwork and the number of paradoxically responding modes in the SST population. The number of unstable modes in the E-PV-VIP subnetwork is quantified by counting the eigenvalues of the Jacobian of the E-PV-VIP subnetwork **J**_*E*−*PV*−*VIP*_ with positive real parts shown on the x-axis. The number of paradoxically responding modes in the SST population is quantified by counting the eigenvalues of the corresponding response matrix **R**_*SS*_ with negative real parts shown on the y-axis. Results for different stimulus sizes are shown in separate panels. Each dot represents one of the ten models. All dots lie along the diagonal lines, indicating that the parity of the number of unstable modes in the E-PV-VIP subnetwork matches the parity of the number of paradoxically responding modes in the SST population. Note that some dots are located at the same coordinates. **B**. Same as A, but for the comparison between the number of unstable models in the E-PV-SST network and the number of paradoxically responding modes in the VIP population. The numbers are quantified by the eigenvalues of the Jacobian of the E-PV-SST subnetwork **J**_*E*−*PV*−*SST*_ with positive real parts shown on the x-axis and the eigenvalues of the corresponding response matrix **R**_*VV*_ with negative real parts shown on the y-axis, respectively. All dots are located at the origin, implying that VIP neurons are not required for stabilization and that patterned perturbations do not elicit paradoxical responses in the VIP population.

**Table S1:** Parameters.

| Symbol | Value | Unit | Description |
| --- | --- | --- | --- |
| $\sigma_{EE}$ | 7 | degree | spatial scale of the Gaussian connectivity profile for E to E connections |
| $\sigma_{EP}$ | 5 | degree | spatial scale of the Gaussian connectivity profile for PV to E connections |
| $\sigma_{ES}$ | 7 | degree | spatial scale of the Gaussian connectivity profile for SST to E connections |
| $\sigma_{EV}$ | 5 | degree | spatial scale of the Gaussian connectivity profile for VIP to E connections |
| $\sigma_{PE}$ | 5 | degree | spatial scale of the Gaussian connectivity profile for E to PV connections |
| $\sigma_{PP}$ | 4 | degree | spatial scale of the Gaussian connectivity profile for PV to PV connections |
| $\sigma_{PS}$ | 5 | degree | spatial scale of the Gaussian connectivity profile for SST to PV connections |
| $\sigma_{PV}$ | 4 | degree | spatial scale of the Gaussian connectivity profile for VIP to PV connections |
| $\sigma_{SE}$ | 7 | degree | spatial scale of the Gaussian connectivity profile for E to SST connections |
| $\sigma_{SP}$ | 5 | degree | spatial scale of the Gaussian connectivity profile for PV to SST connections |
| $\sigma_{SS}$ | 7 | degree | spatial scale of the Gaussian connectivity profile for SST to SST connections |
| $\sigma_{SV}$ | 5 | degree | spatial scale of the Gaussian connectivity profile for VIP to SST connections |
| $\sigma_{VE}$ | 5 | degree | spatial scale of the Gaussian connectivity profile for E to VIP connections |
| $\sigma_{VP}$ | 4 | degree | spatial scale of the Gaussian connectivity profile for PV to VIP connections |
| $\sigma_{VS}$ | 7 | degree | spatial scale of the Gaussian connectivity profile for SST to VIP connections |
| $\sigma_{VV}$ | 4 | degree | spatial scale of the Gaussian connectivity profile for VIP to VIP connections |
| $\sigma_{XL4}$ | 7 | degree | spatial scale of the Gaussian connectivity profile for connections from L4 |
| $\sigma_{XLM}$ | 15 | degree | spatial scale of the Gaussian connectivity profile for connections from LM |
| $\sigma_{XR}$ | 15 | degree | spatial scale of the Gaussian connectivity profile for residual input connections |
| $\sigma_R$ | 15 | degree | spatial scale of the residual input rate field |
| $C_E$ | 1 | a.u. | optimization weighting factor for E responses |
| $C_P$ | 1 | a.u. | optimization weighting factor for PV responses |
| $C_S$ | 1 | a.u. | optimization weighting factor for SST responses |
| $C_V$ | 0.1 | a.u. | optimization weighting factor for VIP responses |

